# Equine piroplasmosis in different geographical areas in France: prevalence heterogeneity of asymptomatic carriers and low genetic diversity of *Theileria equi* and *Babesia caballi*

**DOI:** 10.1101/2024.07.26.605244

**Authors:** M Jouglin, C Bonsergent, N de la Cotte, M Mège, C Bizon, A Couroucé, E Lallemand, L.C. Lemonnier, A Leblond, A Leroux, I Marano, A Muzard, E Quéré, M Toussaint, A Agoulon, L Malandrin

**Affiliations:** INRAE, Oniris, BIOEPAR, 44300, Nantes, France; CISCO-ONIRIS, Department of Equine Internal Medicine, University Veterinary Teaching Hospital, Oniris, 44300 Nantes, France; ENVT, Toulouse, France; VetAgro Sup, Lyon, UMR EpiA, France; ENVA, Maisons-Alfort, France

**Keywords:** : Equine piroplasmosis, asymptomatic carriers, nested PCR, *Theileria equi*, *Babesia caballi*, genotype, 18S rRNA, France

## Abstract

Equine piroplasmosis is a worldwide tick-borne disease caused by the protozoan parasites *Theileria equi* and *Babesia caballi*, with significant economic and sanitary consequences. It can also limit the export of infected horses to piroplasmosis-free countries. These two parasites are genetically variable, with greater diversity observed in *T. equi*. This variability can potentially impact diagnostic accuracy.

Our study aimed to evaluate the frequency of asymptomatic carriers of these parasites in France and describe the circulating genotypes. We used a species-specific nested PCR protocol targeting the 18S small subunit (SSU) rRNA gene and subsequent amplicon sequencing on blood samples collected from 566 asymptomatic horses across four National Veterinary Schools.

The carrier frequency varied considerably, ranging from 18.7% around Paris (central-north) to 56.1% around Lyon (southeast), with an overall prevalence of 38.3%. *T. equi* carriers were ten times more frequent (91.7%, 209/228 isolates) compared to *B. caballi* carriers (8.8%, 19/228 isolates). Notably, *T. equi* carrier frequency was significantly lower in the northern region (Ile de France) compared to the southeastern regions. Interestingly, a strong correlation was observed between the frequencies of asymptomatic carriers and reported cases of acute piroplasmosis across all four geographic areas. Neither gender (female, gelding, or stallion) nor horse age showed a significant effect on the frequency of asymptomatic carriers. In areas with the highest carrier frequency, a substantial proportion of horses (22.2% to 37.5%) carried *T. equi* before the age of three, indicating high infection pressure.

Genotyping of 201 *T. equi* isolates revealed a predominance of genotype E (98%), with only a few isolates belonging to genotype A (2%). Notably, two of the four genotype A isolates were detected in horses originating from Spain. All 19 *B. caballi* isolates belonged to the most common genotype A of this species.

The discussion section explores the link between these results, the tick distribution and abundance, and the frequency of detection of *T. equi* and *B. caballi* in febrile cases attributed to piroplasmosis.

## Introduction

Equine piroplasmosis, a tick-borne disease of equids (horses, donkeys, mules, and zebras), is caused by two protozoan parasites: *Theileria equi* (formerly *Babesia equi*) and *Babesia caballi* (de Waal, 1992; Mehlhorn and Schein, 1998). Over 30 tick species are implicated as possible vectors of equine piroplasmosis (listed in Scoles and Ueti, 2015). However, only species of the genera *Dermacentor*, *Rhipicephalus*, and *Hyalomma* have been confirmed as competent vectors (de Waal, 1992). Equine piroplasmosis has a wide geographical distribution, with its incidence depending on the distribution of these parasite vectors (Onyiche et al., 2019; Tirosh-Levy et al., 2020). Endemic in tropical, subtropical, and some temperate regions, the disease is responsible for significant economic losses to the equine industry due to the treatments and their side effects, reduced animal performance and the negative impact on international trade in horses (Rothschild, 2013).

A few countries, including the United States, Canada, Australia, New Zealand, and Japan, are effectively free of equine piroplasmosis. Consequently, testing for equine piroplasmosis in horses is mandatory, whether for participation in international events or for export to these countries (Knowles, 1996; WOAH, 2023).

Equine piroplasmosis presents with a wide range of clinical signs, from subclinical (apathy, lack of appetite, poor exercise tolerance) to severe (fever, anemia, jaundice, hemoglobinuria, petechial hemorrhages) (Wise et al., 2013; Onyiche et al., 2019; Tirosh-Levy et al., 2020). These symptoms are variable, non-specific, and do not allow differentiation between *B. caballi* or *T. equi* infections. The disease may also be asymptomatic, with infected animals recovering from acute or primary *T. equi* infection remaining carriers for life, even with appropriate treatment. In contrast, horses infected with *B. caballi* may remain carriers for up to four years or be cleared after treatment (Tirosh-Levy et al., 2020).

Clinical signs are therefore unreliable for definitive diagnosis, and carrier status with low parasitemia is challenging for accurate diagnosis. While blood smears can definitively diagnose acute cases, identifying carriers requires more sophisticated techniques. Serological tests like the now-abandoned complement fixation test (CFT) have been developed. However, these methods, including the reliable but time-consuming indirect fluorescent antibody test (IFAT) and the simpler enzyme-linked immunosorbent assay (ELISA), have limitations in sensitivity and specificity due to parasite genetic variability (Wise et al., 2013; Onyiche et al., 2019). The World Organization for Animal Health (WOAH) recommends using IFAT or cELISA for serological diagnosis alongside molecular tests based on parasite DNA amplification (simple, nested, multiplex, RLB, or real-time PCR) (WOAH, 2023). This combined approach offers the most accurate diagnosis for both acute cases and carrier animals.

These tools have also been used to investigate the prevalence of equine piroplasmosis, which is endemic in many parts of Asia, Arabia, South and Central America, Africa, and Europe. Studies have reported widely variable prevalence, reaching up to 88% (reviewed in Onyiche et al., 2020). In Europe, a decreasing south-north gradient of equine piroplasmosis seroprevalence appears to exist, with Italy (Moretti et al., 2010; Piantedosi et al., 2014; Sgorbini et al., 2015; Bartolomé Del Pino et al., 2016), Spain (Camacho et al., 2005; Montes Cortés et al., 2017), Portugal (Ribeiro et al., 2013), and Greece (Kouam et al., 2010) being the most affected. In Ireland (Coultous et al., 2020), the United Kingdom (Coultous et al., 2019), the Netherlands (Butler et al., 2012) and Switzerland (Sigg et al., 2010), the seroprevalence of equine piroplasmosis remains below 5% and a few autochthonous cases are reported in the Netherlands (Butler et al., 2012) and Austria (Dirks et al., 2021). This north-south trend has also been observed across France, with higher serological prevalence in the south (Guidi et al., 2015) compared to the north (Soulé et al., 1990; Le Metayer, 2007; Nadal et al., 2022). The spatial analyses were carried out using the CFT technique, on fairly old data (20 years) and without any knowledge of the clinical status of the animals. Locally high prevalence of equine piroplasmosis in France has been determined on the basis of molecular detection of parasites in geographically limited areas in the South - Camargue, Marseille town and Corsica (Dahmana et al., 2019; Rocafort-Ferrer et al., 2022) - or in geographically unspecified regions (Fritz, 2010).

Molecular tools offer several advantages: they enable sensitive, specific, and reliable detection of pathogens even at low parasite levels (parasitemia) common in carrier horses. Additionally, these tools allow sequencing for studying the genetic variability of isolates, improving our understanding of equine piroplasmosis epidemiology. The genetic diversity of *T. equi* (5 gene clades referred to as A to E) and *B. caballi* (3 clades A, B1 and B2) has been highlighted using the 18S SSU rRNA gene sequences (reviewed in Tirosh-Levy, 2020). Importantly, this diversity can influence the accuracy of serological detection due to the lack of cross-reactions between the antigens used in these tests (Bhoora et al., 2010; Rapoport et al., 2014; Mahmoud et al., 2016). The existing data on equine piroplasmosis prevalence in France suffers from several limitations: reliance on large but outdated data (over 20 years old) collected from a horse population with unspecified status (symptomatic vs. asymptomatic, age, gender) and analyzed with detection tools with low sensitivity/specificity (CFT) that are no longer used (Le Metayer, 2007; Nadal et al., 2022), or on geographically limited horse populations (Dahmana et al., 2019; Rocafort-Ferrer et al., 2022). Our study aimed to address these gaps. We employed molecular tools to investigate the carriage of *T. equi* and *B. caballi* in asymptomatic horses across four large geographically distinct regions of France. We focused on asymptomatic horses because they represent a significant parasite reservoir, particularly for *T. equi*. Additionally, we assessed the genetic diversity of these two pathogens, as this variation can impact diagnostic results.

## 2. Materials and methods

### 2.1. Animals and sampled areas

Blood samples were collected from equids presented at four French Veterinary Schools in Maisons-Alfort (ENVA, north-central), Nantes (Oniris, west), Lyon (VetAgro Sup, southeast) and Toulouse (ENVT, southwest). Each school was considered as a focal point with horses located in their surroundings. For clarity in tables and figures, the veterinary schools will be referred to by the main city names: Nantes (Oniris), Paris (ENVA), Toulouse (ENVT) and Lyon (VetAgro Sup).

The study was presented to and approved by the Oniris Ethics Committee for Veterinary Clinical and Epidemiological Research (CERVO-2019-1-V). After explaining the purpose of the study, written consent was obtained from each horse owner for their animal’s participation. During sample collection, owners completed questionnaires providing information about their horses’ characteristics, including location at sampling time, breed, gender, age, previous geographic locations, and history of acute piroplasmosis.

The horses recruited were presented at veterinary schools for various reasons (surgery, reproduction, lameness, etc.) but none related to equine piroplasmosis symptoms. Blood was collected by jugular venipuncture into sterile heparinized tubes. Heparin was chosen as the anticoagulant because it preserves parasite viability for up to 9 days at room temperature, ensuring good DNA quality for analysis (Malandrin et al., 2004). The blood samples and questionnaires were sent to the BIOEPAR research unit in Nantes for analysis.

As this study was part of the national citizen science program PiroGoTick, the results were made anonymous and accessible to participants on the project website https://www.pirogotick.fr. Each participant received an information note explaining the project and how to retrieve the results using their unique reference on the PiroGoTick website.

### 2.2 Generation of geographical location maps of horses

For each of the four veterinary schools (Paris, Toulouse, Lyon, and Nantes), we generated geographical location maps of the horses included in this study. These maps were constructed using data from the French National Geographic Institute (IGN) (including GPS location of communes and a map of mainland France in Lambert93 format) and information on horse origin collected through questionnaires at admission time. All data were then summarized into maps using QGIS software (v3.36).

### 2.3. Molecular detection of blood piroplasms

Genomic DNA was extracted from 200 μL of pelleted red blood cells diluted 1:1 in PBS1X using the DNA Nucleospin blood extraction kit following the manufacturer’s instructions (Macherey-Nagel). The extracted DNA was then stored at -20°C for later use.

To detect the presence of piroplasms, we first amplified a region of the 18S rRNA gene, a common genetic marker for these parasites, using a standard PCR technique and the primers CRYPTOF and CRYPTOR (Malandrin et al., 2010). Reactions were carried out in 30 μL reaction mixtures containing 1 X buffer, 4 mM MgCl2, 0.2 mM of each dNTP (Eurobio), 1 unit GoTaq G2 Flexi DNA Polymerase (Promega), 1 μM of each primer and 7 μL of DNA template. PCR cycling comprised 5 min at 95◦C, 40 cycles at 95◦C for 30 s, 60 s at 58°C, 30 s at 72°C, and a final extension at 72◦C for 5 min. Nested PCRs were then performed to specifically and separately amplify *B. caballi* and *T. equi*, with primers specific for *B. caballi* (BabF4 and BabR4, expected amplicons length of 856 bp) and *T. equi* (TeqF3 and TeqR5, expected amplicons length of 1470 bp). We used 10 μL of 1/100 diluted amplicons in a 30 μL reaction mixture containing the same components as the first PCR. Cycling conditions were the same as for the primary reaction, with the exception of the annealing temperature, which was 54°C in both cases of second specific amplification. The primers specific to *B. caballi* (BabF4 5’-TTGTAATTGGAATGATGGC-3’ and BabR4 5’-TCCCTACAACTTYTCGRTGT-3’) and the primers specific to *T. equi* (TeqF3 5’-TCAGTTGCGTTTATTAGAC-3’ and TeqR5 5’-CACAAAACTTCCCTAGACG-3’) were designed for the present study based on alignments of NCBI sequences selected to represent the different genetic clades.

Amplified fragments were purified with ExoSAP-IT reagent following the manufacturer’s instructions (Affymetrix). Mono or bi-directional Sanger sequencing was then performed by Eurofins Genomics, Germany, using the original PCR primers. The resulting sequences were assembled using Geneious R6 software (https://www.geneious.com). To identify sequence similarities, an online BLAST search was conducted (http://blast.ncbi.nlm.nih.gov). For further analysis, ClustalW software (https://www.ebi.ac.uk) was used to perform sequence similarity analysis. Finally, representative sequences were deposited in GenBank (accession numbers PQ044745 to PQ044786 for T. equi; accession numbers PQ044787 to PQ044795 for B. caballi).

### 2.4. Statistical analysis

Data analysis was conducted in R (v4.3.3). The normality of all continuous variables was tested using the Shapiro-Wilk normality test. Continuous variables were expressed as mean, median, interquartile range (IQR) and range. Differences in the proportional distribution of categorical variables among two groups were evaluated using Fisher’s exact tests. Differences in the distribution of continuous variables between two groups were evaluated using the Wilcoxon rank sum test. For multiple pairwise comparisons of categorical or continuous variables, a Bonferroni correction was applied: the significance level α was set to 0.05/n, with n the number of comparisons.

### 2.5. Phylogenetic Analysis

To construct maximum likelihood phylogenetic trees, the advanced PhyML+SMS/OneClick workflow available on the NGPhylogeny.fr website was employed (https://ngphylogeny.fr/). In brief, this pipeline performs several steps: first, it aligns sequences in FASTA format using MAFFT software (Katoh and Standley, 2013). Then, it cleans the alignments using BMGE software (Criscuolo and Gribaldo, 2010). Finally, it constructs phylogenetic trees using PhyML software (Guindon et al., 2010). This process includes automatic selection of the most suitable evolutionary model using SMS software (Lefort et al., 2017) and generates the final tree in Newick format (Junier and Zdobnov, 2010). The parameters used are the default ones except for the PhyML+SMS analysis where the following choices were made: use of the BIC (Bayesian Information Criterion) for the model selection criterion, the NNI (Nearest Neighbor Interchange) for the tree topology and a boostrapping (FTB+TBE) of 1000 replicates for inference (Lemoine et al., 2018).

Phylogenetic analyses of the sequences using Bayesian inference were performed using MAFFT software (Katoh and Standley, 2013), followed by cleaning of these alignments using BMGE software (Criscuolo and Gribaldo, 2010). We then used the BEAST2 software (v2.7.6) (Bouckaert et al.,2019) with the bModelTest plugin (v1.3.3) (Bouckaert and Drummond, 2017) to build the trees.

The generated trees were then formatted using TreeViewer v 2.2.0 (Bianchini and Sánchez-Baracaldo, 2024).

## 3. Results

### 3.1. Description of sampled equids’ population

From November 2019 to January 2023, blood samples from 566 horses were collected and analyzed, all asymptomatic for piroplasmosis. The place of residence of each horse is shown in Figure 1 to highlight the geographical distribution of the equine population studied.

**Fig. 1.**
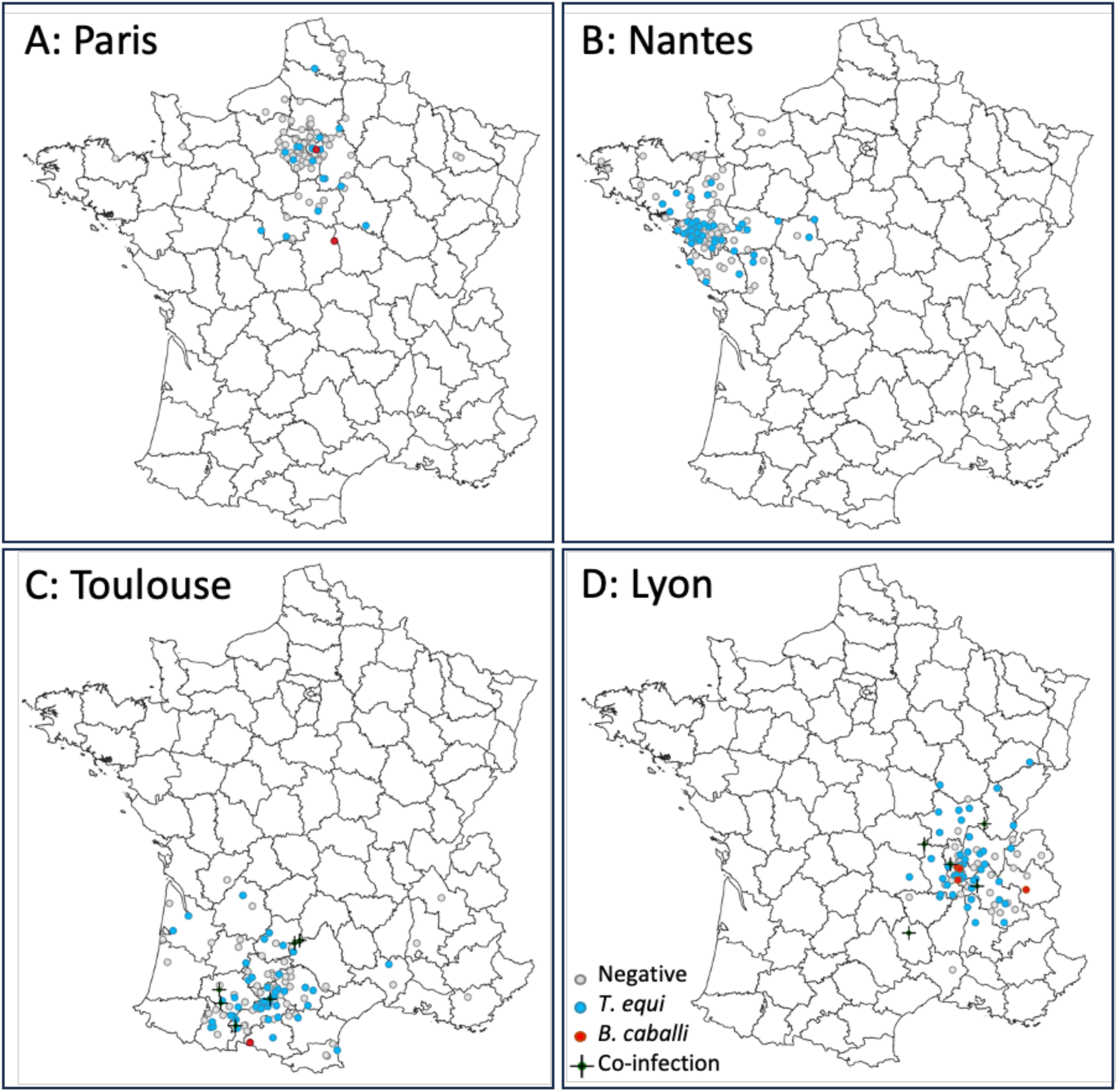
Geographicall location and carrier status of the horse population samplled in the four veterinary schools in France between 2019-2023. A: 107 horses around Paris. B: 169 horses around Nantes. C: 135 horses around Toulouse. D: 155 horses around Lyon. The trench departments borders are indicated.

The study in Nantes ran from 2019 to 2023 and enrolled 169 horses from 11 counties (Table 1). At the other veterinary schools, blood was collected between 2020-2022 from 107 equids located in 22 different counties around Paris, 135 equids from 20 counties around Toulouse and 155 equids from 16 counties around Lyon (Table 1). In total, equids from 62 different counties were sampled, with equids from the same county being presented at two different veterinary schools (Nantes/Paris and Lyon/Toulouse, Fig. 1). A highly variable number of equids were sampled in these counties (1 to 100). A detailed breakdown of the equine population studied by sampling year and department is provided in supplementary tables 1 and 2, respectively.

**Table 1.** Description of the sampled horse population (sampled area, year of sampling and gender).

| Veterinary School | Number of tested equids per year |  |  |  |  | Sex |  |  |  | Total |
| --- | --- | --- | --- | --- | --- | --- | --- | --- | --- | --- |
|  | 2019 | 2020 | 2021 | 2022 | 2023 | Female | Gelding | Stallion | ND |  |
| Paris | 0 | 39 | 53 | 15 | 0 | 40 | 61 | 6 | 0 | 107 |
| Nantes | 71 | 62 | 32 | 0 | 4 | 73 | 74 | 22 | 0 | 169 |
| Toulouse | 0 | 5 | 52 | 78 | 0 | 57 | 64 | 12 | 2 | 135 |
| Lyon | 0 | 18 | 87 | 50 | 0 | 80 | 50 | 15 | 10 | 155 |
| Total | 71 | 124 | 224 | 143 | 4 | 250 | 249 | 55 | 12 | 566 |

Incomplete questionnaires from some horse owners resulted in missing data for about 2% of the population. The sex distribution of the horses sampled across the four veterinary schools differed. Overall, the study population consisted of a similar proportion of females (250) and geldings (249), with approximately 10% being males (Table 1). Notably, the ratio of geldings to females was balanced in Nantes and Toulouse, but skewed towards females in Lyon and geldings in Paris. A chi-squared test revealed a statistically significant difference (p<0.05) in sex distribution among the four sampled populations.

(Albert fichier pdf Pearson’s Chi-squared test)

The average age of the horse population studied was 11.5 years, ranging from 1 to 31 years. Horses in Nantes were significantly younger (mean 10.4 years, range 1-31 years) compared to the other veterinary schools, which showed similar age distributions (average 11.8 years, range 2-26 years in Paris; 11.9 years, range 1-31 years in Toulouse; 12 years, range 1-27 years in Lyon). A violin diagram illustrating the age distribution across the veterinary schools is provided in supplementary Fig. 1.

### 3.2. Frequency of asymptomatic piroplasmosis carriers according to the geographical location

Our study revealed a carrier frequency of 38.3% (217 out of 566 horses) for equine piroplasmosis, with *T. equi* being the predominant parasite (91.7%; 209/228 isolates). *B. caballi* was less frequent (8.3%; 19/228 isolates) (Table 2). A small proportion of horses (1.9%) was co-infected with both parasites. Over half (57.8%) of *B. caballi* carriers were co-infected with *T. equi*. Single infections with *B. caballi* were less common (1.4%, 8 horses). Most *B. caballi* carriers were identified in the south of France, around Toulouse and Lyon (89.5%), with none found in Nantes and 10.5% identified around Paris.

**Table 2.** Sampled population and prevalence of asymptomatic carriers of piroplasmosis, *T. equi* and *B. caballi* in equids sampled in the 4 veterinary schools between 2019 and 2023.

| Veterinary School | Period | Number of sampled equids | Carrier frequency % [95% CI] (number) of |  |  |  |
| --- | --- | --- | --- | --- | --- | --- |
|  |  |  | Piroplasmosis carriers | *T. equi* carriers | *B. caballi* carriers | Co-infected carriers |
| Paris | 2020-2022 | 107 | 18.7 [12.1-27.6] (20) | 16.8 [10.5-25.6] (18) | 1.9 [0.3-7.2] (2) | 0 [0.0-4.3] (0) |
| Nantes | 2019-2023 | 169 | 32.0 [25.1-39.6] (54) | 32.0 [25.1-39.6] (54) | 0 [0.0-2.8] (0) | 0 [0.0-2.8] (0) |
| Toulouse | 2020-2022 | 135 | 41.5 [33.2-50.3] (56) | 40.7 [32.5-49.5] (55) | 5.2 [2.3-10.8] (7) | 4.4 [1.8-9.8] (6) |
| Lyon | 2020-2022 | 155 | 56.1 [47.9-64.0] (87) | 52.9 [44.8-60.9] (82) | 6.5 [3.3-11.9] (10) | 3.2 [1.2-7.8] (5) |
| Total | 2019-2023 | 566 | 38.3 [34.3-42.5] (217) | 36.9 [33.0-41.1] (209) | 3.4 [2.1-5.3] (19) | 1.9 [1.0-3.6] (11) |

The frequency of equine piroplasmosis carriers varied considerably between the four horse populations sampled: 18.7% around Paris, 32.0% around Nantes, 41.5% around Toulouse and 56.1% around Lyon (Table 2). *T. equi* carrier frequencies were significantly lower around Paris compared to the other three populations and significantly higher in Lyon compared to the two northern populations (Nantes and Paris) (Table 3). The frequencies of equine piroplasmosis in Nantes and Toulouse were not statistically significantly different, were those between Toulouse and Lyon.

**Table 3.**
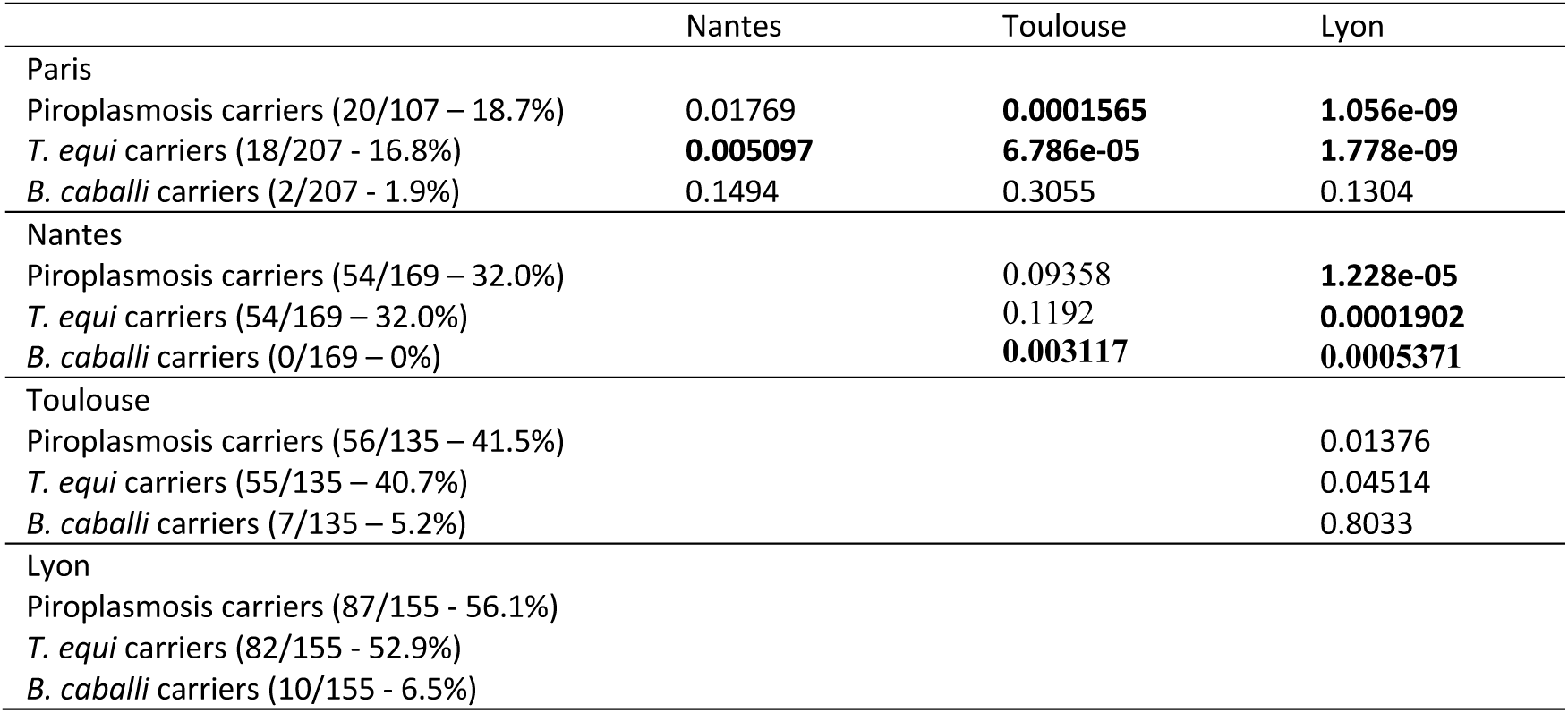
Statistical analysis of carrier frequencies between the 4 sampled populations. p-values of 2 by 2 frequency comparisons of carriers using the Fisher’s exact test are indicated. The significance threshold was set at 0.0083 after the Bonferroni correction and the significant differences are highlighted in bold.

|  | Nantes | Toulouse | Lyon |
| --- | --- | --- | --- |
| Paris |  |  |  |
| Piroplasmosis carriers (20/107 – 18.7%) | 0.01769 | **0.0001565** | **1.056e-09** |
| *T. equi* carriers (18/207 - 16.8%) | **0.005097** | **6.786e-05** | **1.778e-09** |
| *B. caballi* carriers (2/207 - 1.9%) | 0.1494 | 0.3055 | 0.1304 |
| Nantes |  |  |  |
| Piroplasmosis carriers (54/169 – 32.0%) |  | 0.09358 | **1.228e-05** |
| *T. equi* carriers (54/169 – 32.0%) |  | 0.1192 | **0.0001902** |
| *B. caballi* carriers (0/169 – 0%) |  | **0.003117** | **0.0005371** |
| Toulouse |  |  |  |
| Piroplasmosis carriers (56/135 – 41.5%) |  |  | 0.01376 |
| *T. equi* carriers (55/135 – 40.7%) |  |  | 0.04514 |
| *B. caballi* carriers (7/135 – 5.2%) |  |  | 0.8033 |
| Lyon |  |  |  |
| Piroplasmosis carriers (87/155 - 56.1%) |  |  |  |
| *T. equi* carriers (82/155 - 52.9%) |  |  |  |
| *B. caballi* carriers (10/155 - 6.5%) |  |  |  |

Despite the low overall frequency of *B. caballi* carriers, equids in the areas around Toulouse and Lyon had a significantly higher frequency compared to Nantes alone, but not compared to Paris. There was no statistical difference between Lyon and Toulouse (Table 3).

More detailed geographical representations of the horse location and carrier frequency by French region are shown in Fig. 1 and 2, respectively. Data for Normandy, Grand-Est, Provence-Alpes-Côte d’Azur, and Corsica were excluded due to limited samples (less than 10 horses) (see Supplementary Table 2 for details). The frequency of equine piroplasmosis carriers was significantly higher in Burgundy-Franche-Comté (82.4%) compared to the five western or northern regions, but not compared to the southern or neighboring regions (Supplementary table 3). The Ile-de-France region had a significantly lower carrier frequency compared to the southeastern regions: Burgundy-Franche-Comté, Auvergne-Rhône-Alpes, and Occitanie.

**Fig. 2.**
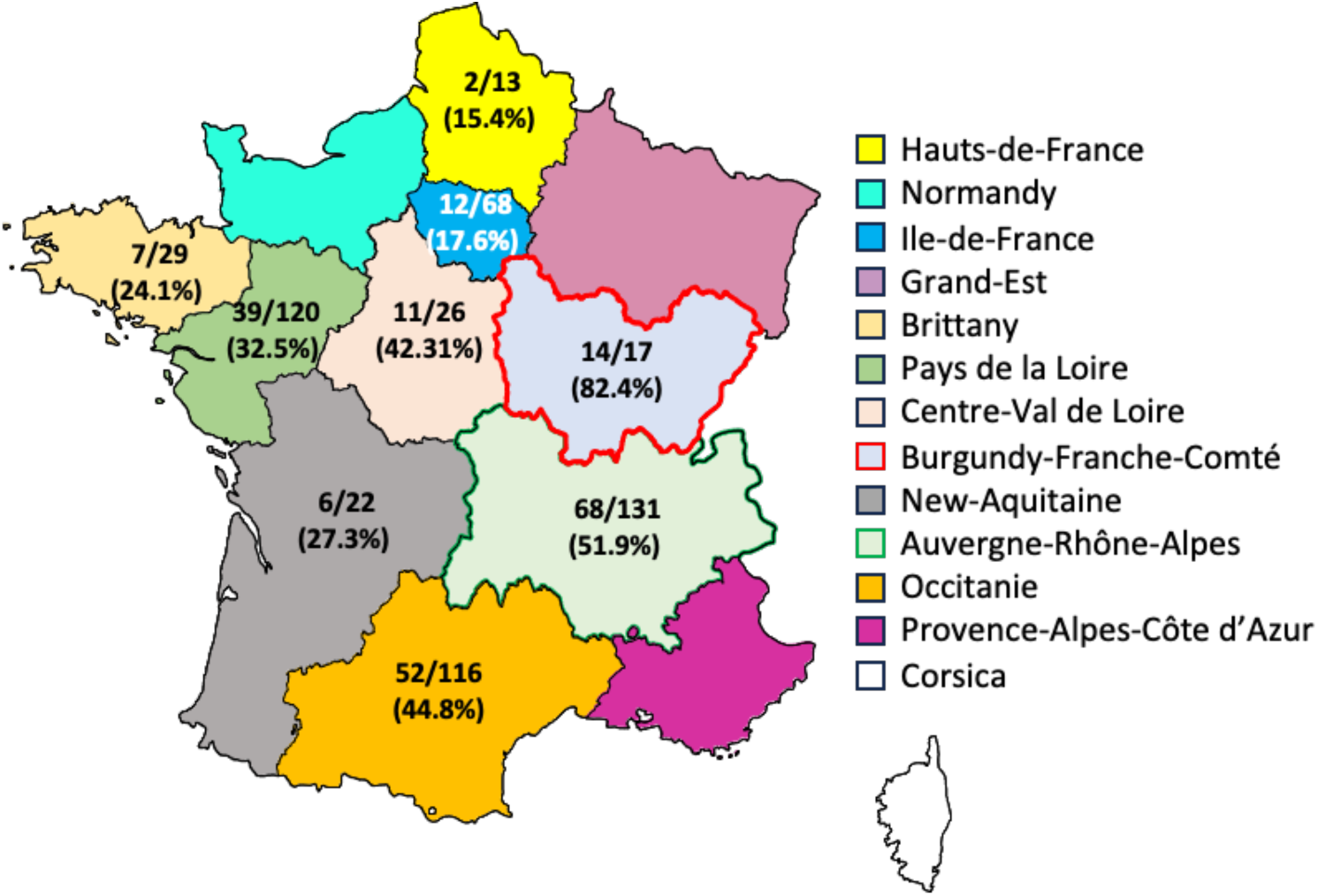
Frequency of equine piroplasmosis carriers in the regions of France. The numbers of positive and tested horses are indicated in each region as well as the deduced frequency. Data are not indicated in the regions Normandy, Grand-Est, PCA and Corsica due to the too low number or abence of sampled equids (respectively 8, 2, 1 and 0). The regions where the frequency of carriers is significalty higher in France are indicated with a red border (Burgundy-Franche-Comte) and a green border (Auvergne­ Rhone-Alpes).

### 3.3. Comparison of frequencies of asymptomatic and symptomatic horses in the sampled population

Questionnaires asked horse owners about any previous episode of acute equine piroplasmosis in their horses, including the diagnostic methods used and the parasite species identified. Owners were more certain about past clinical signs of piroplasmosis than the specific methods or parasite species involved. The reported frequencies of previous acute piroplasmosis cases were lower but correlated positively with the carrier frequencies observed in this study (Fig. 3). According to the questionnaires, diagnoses during acute phases identified similar proportions of *T. equi* (21 cases) and *B. caballi* (17 cases) as the presumed causative agent. We did not attempt a case-by-case study for reasons detailed in the discussion section.

**Fig. 3.**
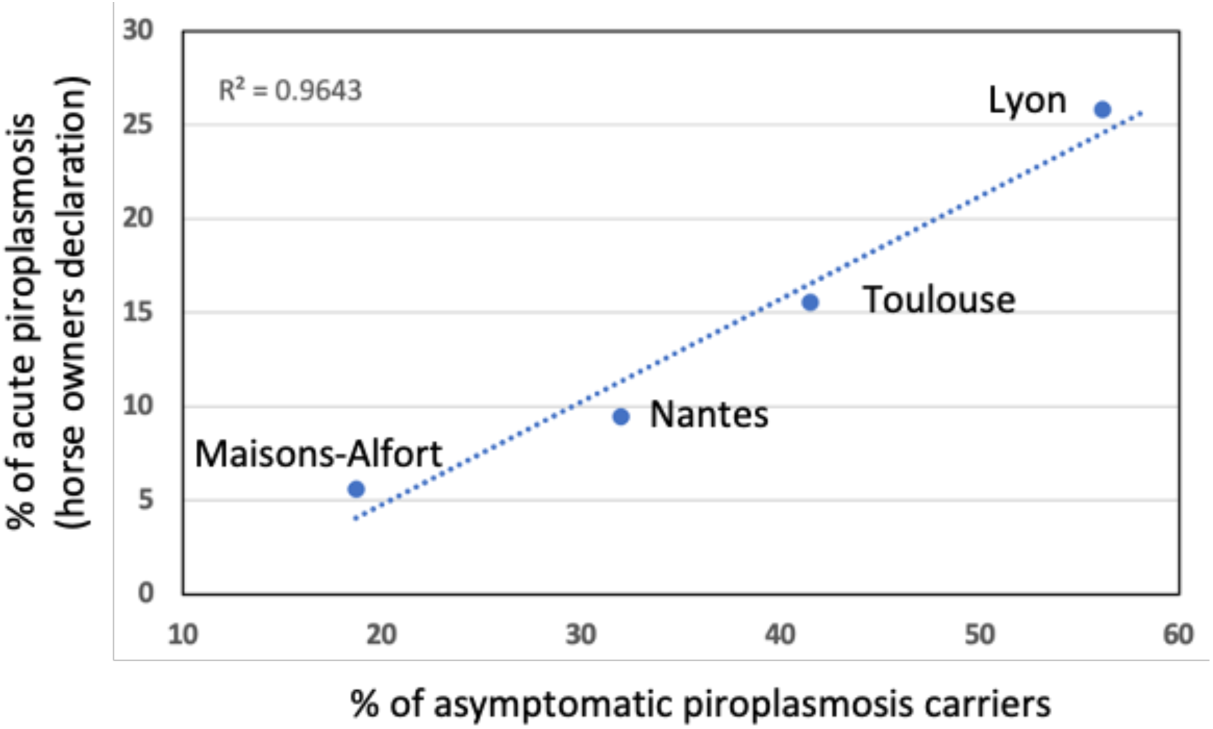
Correlation between the frequency of carriers of equine piroplasmosis and the percentage of horses with reported acute piroplasmosis in the past, in each of the 4 equine populations analyzed around Paris, Nantes, Toulouse and Lyon. Acute piroplasmosis occurence is based on owners declarations in the questionnaires.

### 3.4. Influence of horse movement or origins on piroplasmosis carrier status

The aim of this section was not to analyze horse movements in detail, but to highlight how they can influence the estimated frequency of carrier status, particularly in regions of low endemicity. We therefore chose Paris as our case study. Among the 20 carrier horses, nine (45%) were born or lived in regions where equine piroplasmosis is highly endemic: five from French regions (Jura, Yonne, Burgundy, Toulouse, Corsica) and four from southern countries (Italy, Spain). In contrast, only 13 of the 87 non-carrier horses (15%) originated from these same regions or countries. Originating from a region or country with high piroplasmosis frequency was associated with a statistically significantly higher risk of infection (p=0.005, Fisher’s exact test). This highlights how horse movement can potentially inflate carrier frequency estimates in areas with low endemicity, such as the Paris region.

### 3.5. Influence of age and gender on the piroplasmosis carrier status

Due to differences in carrier frequencies observed between regions and between parasite species, we performed separate statistical analyses for carrier frequency according to age groups for each region. Additionally, analyses were performed separately for *T. equi* and *B. caballi* carriers due to their distinct infection dynamics. *T. equi* persists for life, potentially leading to a higher carrier prevalence with age. Conversely, *B. caballi* infections can be cleared, with a possible decrease in carrier frequency among older horses due to acquired immunity.

The frequencies of *T. equi* and *B. caballi* carriers by age group for each collection site are detailed in Table 4.

**Table 4.** Frequencies of piroplasmosis carriers (*T. equi* and *B. caballi*) by age classes in the 4 sampled horse populations.

|  | *T. equi* carrier frequency (%) |  |  |  | *B. caballi* carrier frequency (%) |  |  |
| --- | --- | --- | --- | --- | --- | --- | --- |
|  | Paris | Nantes | Toulouse | Lyon | Paris | Toulouse | Lyon |
| <= 2 Y | 0.0 | 27.8 | 22.2 | 37.5 | 0.0 | 11.1 | 25 |
| <= 5 Y | 11.1 | 20.0 | 28.1 | 44.4 | 0.0 | 9.4 | 13.9 |
| 6 to 10 Y | 16.2 | 33.3 | 44.8 | 65.5 | 5.4 | 5.3 | 3.5 |
| 11 to 15 Y | 18.9 | 31.0 | 40.8 | 51.2 | 0.0 | 0.0 | 2.3 |
| 16 to 20 Y | 23.5 | 43.5 | 40.0 | 53.6 | 0.0 | 10.0 | 10.7 |
| >20 Y | 0.0 | 44.4 | 42.9 | 56.3 | 0.0 | 0.0 | 0.0 |

We analyzed carrier frequency in specific age groups (e.g., 0-5 years, 6-10 years, etc.). In all four sampled populations, the frequencies of *T. equi* carriers were high for horses of 5 years or less, with up to 44.4% of horses already carrying *T. equi* around Lyon. More than ⅓ of the horses were already infected before the age of 3 years around Lyon, and about ¼ around Nantes and Toulouse. In contrast to *T. equi*, the pattern for *B. caballi* carriers differed. Around Lyon and Toulouse, a significant proportion of carriers were young (less than 3 years old): 40% (4 out of 10) around Lyon and 42.9% (3 out of 7) around Toulouse. Interestingly, these regions also showed a notable presence of *B. caballi* carriers in the 16-20 year age class.

The observed gradient in *T. equi* carrier frequency across the four regions (Paris < Nantes < Toulouse < Lyon) remained consistent for most age groups, except for the oldest horses. In the southern populations (Toulouse and Lyon), the carrier frequency appeared to plateau after 10 years of age, while it continued to rise in the northern regions (Paris and Nantes). However, applying a Bonferroni correction to adjust the significance level (p > 0.001), no statistically significant effect of age on the frequencies of either *T. equi* or *B. caballi* carriers was detected within any age group (detailed data not shown).

The population of sampled males was significantly younger than that of females or geldings in all four sampled areas (Table 5). Despite this age difference, piroplasmosis carrier frequencies were not significantly different between stallions (36.4%; 20/55) and geldings (35.7%; 89/249) (p=1) or females (40.4%; 101/250) (p=0.65), nor between females and geldings (p=0.31, Fisher’s exact test). This analysis was performed for each sampled area separately with the same results.

**Table 5.** Piroplasmosis carrier frequencies and ages of the horse population sampled in the 4 sampled areas according to the gender and infectious status.

| Frequency of piroplasmosis carriers (%) and mean age (min/max) of each category |  |  |  |  |  |  |  |  |
| --- | --- | --- | --- | --- | --- | --- | --- | --- |
| Sampled area | Studied population |  | Female |  | Gelding |  | Stallion |  |
|  | Carrier freq. | Age total<br>*Age Healthy*<br>**Age Infected** | Carrier freq. | Age total<br>*Age Healthy*<br>**Age Infected** | Carrier freq. | Age total<br>*Age Healthy*<br>**Age Infected** | Carrier freq. | Age total<br>*Age Healthy*<br>**Age Infected** |
| Paris | **18.7%** | 11.8 (2-26)<br>*11.9 (2-26)*<br>**11.6 (5-18)** | **17.5%** | 11.2 (4-23)<br>*11.2 (4-23)*<br>**11.1 (5-16)** | **21.3%** | 12.5 (2-26)<br>*12.6 (2-26)*<br>**11.9 (7-18)** | **0%** | 9.2 (3-19)<br>*9.2 (3-19)*<br>**nd** |
| Nantes | **32.0%** | 10.4 (1-31)<br>*9.6 (1-31)*<br>**12.1 (1-24)** | **26.0%** | 10.4 (1-26)<br>*9.8 (1-26)*<br>**12.1 (2-24)** | **36.5%** | 11.5 (2-31)<br>*10.4 (2-31)*<br>**13.5 (2-23)** | **36.4%** | 6.0 (1-24)<br>*5.7 (1-24)*<br>**6.6 (1-16)** |
| Toulouse | **41.5%** | 11.9 (1-31)<br>*11.6 (1-28)*<br>**12.4 (1-31)** | **45.6%** | 13.3 (1-31)<br>*13.5 (1-28)*<br>**13.1 (1-31)** | **37.5%** | 12.3 (1-25)<br>*11.6 (1-25)*<br>**13.5 (6-25)** | **41.7%** | 3.3 (1-10)<br>*3.3 (1-10)*<br>**3.2 (1-6)** |
| Lyon | **56.1%** | 12.0 (1-27)<br>*11.4 (1-23)*<br>**12.6 (1-27)** | **61.3%** | 13.4 (2-27)<br>*13.2 (2-23)*<br>**13.5 (4-27)** | **50.0%** | 12.4 (2-25)<br>*11.5 (2-23)*<br>**13.5 (2-25)** | **46.7%** | 3.6 (1-14)<br>*3.4 (1-7)*<br>**3.9 (1-14)** |
| Total population | **38.3%** | 11.5 (1-31)<br>*11.0 (1-28)*<br>**12.3 (1-31)** | **40.4%** | 12.2 (1-31)<br>*11.6 (1-28)*<br>**13.0 (1-31)** | **35.7%** | 12.1 (1-31)<br>*11.5 (1-26)*<br>**13.3 (2-25)** | **36.4%** | 5.1 (1-24)<br>*5.3 (1-24)*<br>**4.7 (1-16)** |

### 3. Genetic diversity of *T. equi* and *B. caballi* and phylogenetic analysis

In total, 201 samples produced 18S rRNA sequences identified as *T. equi*. These sequences were from horses in the areas around Paris (18), Nantes (52), Toulouse (53) and Lyon (78). Sequence lengths ranged between 230 and 1412 bp. Sequence identities were obtained via a blast analysis with reference sequences selected to represent geographically distant locations (AY534882 from Spain and KF559357 from China for the genotype E, KJ573370 from Brazil, AY150062 from Spain and KX227625 from Israel for the genotype A). Isolates and sequences description are detailed in the Supplementary tables 4 and 5. A majority of genotype E (197/201, 98%) and a few sequences of genotype A (4/201, 2%) were characterized.

Within clade E, the sequence identities were higher than 99% identity to each other or with geographically distant genotype E isolates, while identities were lower than 98.5% with genotype A reference sequences. The genotype E isolates originated from equids living in 43 different counties in France.

The 4 sequences belonging to *T. equi* genotype A demonstrated identities higher than 99% to each other and to geographically distant reference sequences. Identities with genotype E sequences were below 97%. The genotype A isolates originated from equids living in 3 different counties in France. Two of these horses originated from Spain.

All 19 sequences identified as *B. caballi* belonged to the genotype A, and they shared identities higher than 99% to each other and to sequences of *B. caballi* genotype A from geographically distant countries (Supplementary table 6).

Representative sequences of *T. equi* genotype E (38 sequences from different counties with accession numbers PQ044745 to PQ044782), *T. equi* genotype A (4 sequences with accession numbers PQ044783 to PQ044786) and *B. caballi* (9 sequences from different counties with accession numbers PQ044787 to PQ044795) were deposited in GenBank. Details on sequence lengths, identities and geographic locations are given in the supplementary tables 4 to 6.

Representative sequences for each *T. equi* genotype and for *B. caballi* evidenced in this study were selected to confirm their phylogenetic positioning, with either the Maximum likelihood method (Fig. 4 and 5) or with the bayesian inference method (Supplementary Fig. 2 and 3). Positioning in the different genotype clades was identical for each sequence whatever the phylogenetic method used.

**Fig. 4.**
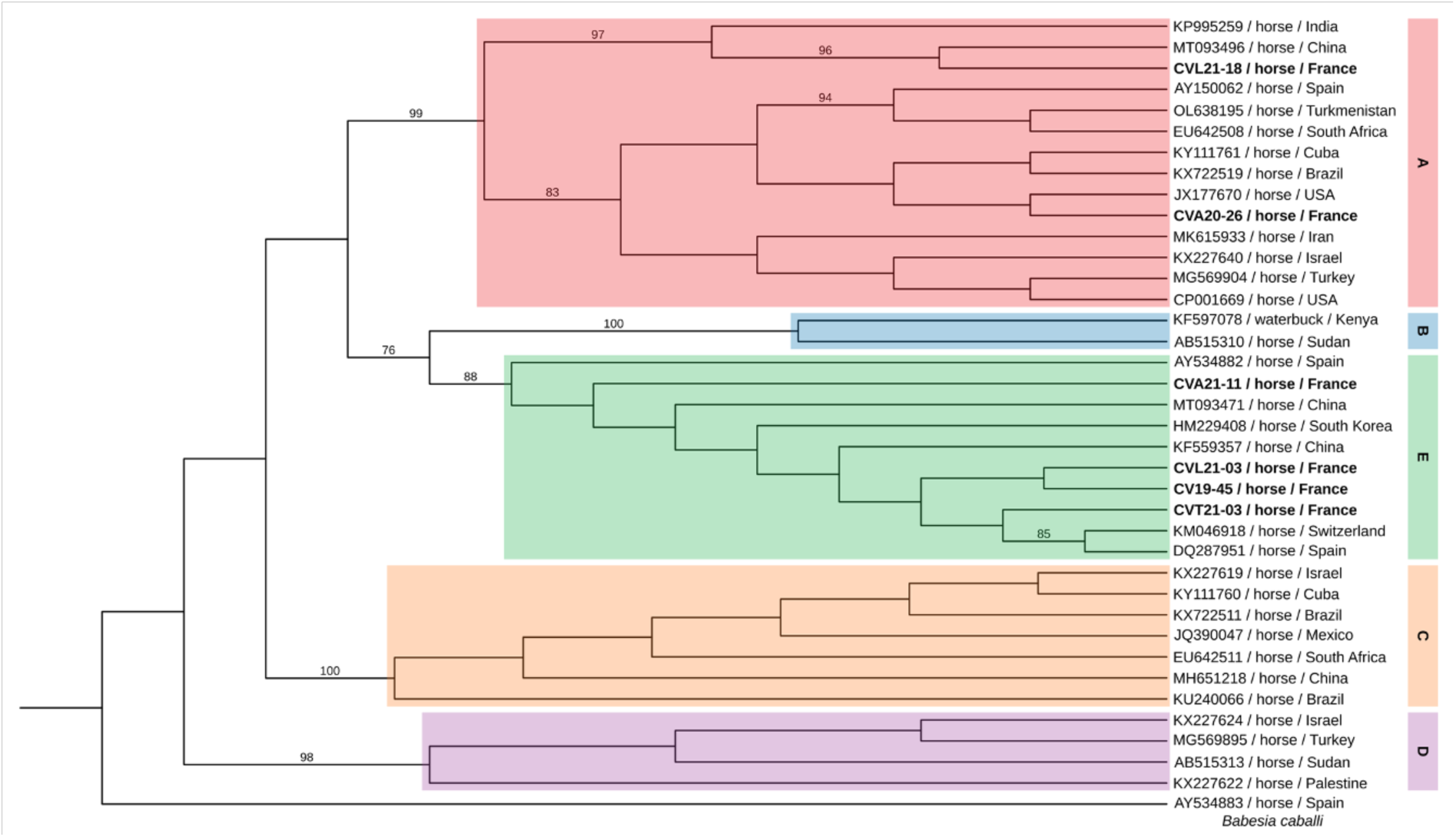
A maximum likelihood tree of *Theileria equi* isolates based on sequences derived from the amplified 18S rRNA hypervariable regions. The tree includes 38 sequences with 1295 analyzed positions. The Tamura-Nei model has been used with 1000 repetitions. Significant bootstrap values (>70) are indicated on the branches. *T. equi* sequences reported in the litterature are denoted by their GenBank accession number, host and country of origin. Representative sequences obtained in the present study are indicated in bold. Previously desribed genotypes (A-E) as described in Tirosh­ Levy et al. in 2020 are highlighted.

**Fig. 5.**
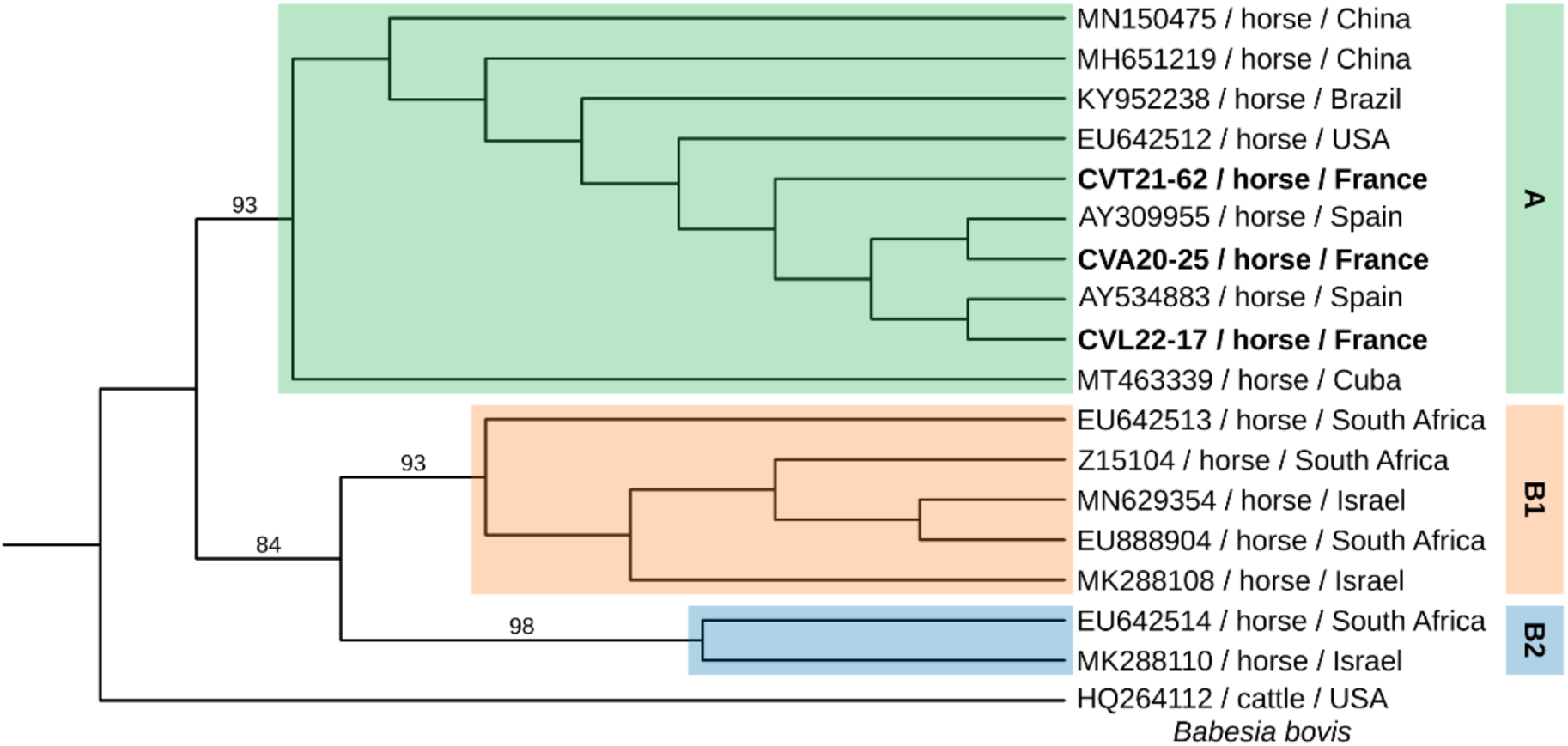
A maximum likelihood tree of *Babesia cabal/i* isolates based on sequences derived from the amplified 18S rRNA hypervariable regions. The tree includes 18 sequences with 860 analyzed positions. The Tamura-Nei model has been used with 1000 repetitions. Significant bootstrap values (>70) are indicated on the branches. *B. cabol/i* sequences reported in the litterature are denoted by their GenBank accession number, host and country of origin. Representative sequences obtained in the present study are indicated in bold. Previously desribed genotypes (A, Bl and 82) as described in Tirosh-Levy et al. in 2020 are highlighted.

## 4. Discussion

The aim of this study was to investigate the prevalence of asymptomatic carriers of equine piroplasmosis in France, analyze the relative proportion of each causative agent (*T. equi* and *B. caballi*), and to genetically characterize these parasite species. One of the inclusion criteria for horses in this study was outdoor or mixed housing (mainly outdoor pasture/indoor), in order to focus on animals in contact with ticks, potential vectors. Blood samples were taken from horses in four areas around each national veterinary school in France.

In the four regions studied, the frequency of *T. equi* carriers was significantly higher (16.8-52.9%) than that of *B. caballi* (0-6.5%), and *B. caballi* could not be evidenced in asymptomatic horses from the North-western part of France (Nantes). This pattern aligns with two other studies involving asymptomatic horses in southern France (Rocafort-Ferrer et al., 2022; Dahmana et al., 2019), and most other European countries where equine piroplasmosis is endemic (Montes Cortés et al., 2016; Nadal et al., 2022). This occurs despite the presence of vector ticks common to both piroplasm species in these countries. It probably reflects the fact that infected animals that survive a first infection completely eliminate *B. caballi* 1 to 4 years after infection or after appropriate treatment, whereas *T. equi* remains as a lifelong infection (de Waal and Van Heerden, 1994).

We observed variations in the frequencies of equine piroplasmosis across different regions in France. The highest carrier frequencies were noted in the two southern areas studied, particularly around Lyon (56.1%), An intermediate frequency was recorded around Nantes (32.0%), and a lower frequency was found around Paris (18.7%), in the central-northern part of France. A significantly higher prevalence was measured in the Burgundy-Franche-Comté (82.4%), and high prevalence was also found in Auvergne-Rhône-Alpes (51.9%) and Occitanie (44.8%). This apparent gradient in the prevalence of piroplasmosis, increasing from the north-west to the south-east, has also been documented using serological data based on the Complement Fixation Test (CFT) at the scale of French departments (Le Metayer, 2007), and later re-analyzed at the regional level (Nadal et al., 2022), using data from an impressive dataset of 16,127 horse sera. According to these analyses, the regions of Burgundy-Franche-Comté, Auvergne-Rhône-Alpes and Occitanie also exhibited the highest percentages of seropositivity. However, the seroprevalence measured 20 years ago in these regions was low (maximum 30%) compared to the prevalence of carriers determined in the present study. These differences could be explained by an increase in the prevalence of equine piroplasmosis and/or the sensitivity/specificity of the method used. In the southernmost Mediterranean part of France, equine piroplasmosis agents were detected in the blood of 75% of horses from stables where unexplained fever or weight loss had been identified by local veterinarians (Rocafort-Ferrer et al., 2022). In the same area, using the same kind of method (real-time PCR), lower prevalence was measured (29.4%) in horses from equestrian centers, where access to pasture and therefore to vector ticks may have been more restricted (Dahmana et al., 2019). Differences in the prevalence of equine piroplasmosis between horses with similar types of housing (pasture in the present study) but in different geographical locations could be related to different climatic conditions. These conditions would influence not only the presence of different vector tick species, but also their abundance and activity period length (Kouam et al., 2010; Grandi et al., 2011; García-Bocanegra et al., 2013). Climatic factors such as temperature, humidity or rainfall influence tick habitats, and most likely the transmission dynamics of equine piroplasmosis. In this study, the four sampling areas are subject to different climates (Joly et al., 2010). Nantes experiences an oceanic climate, while the climate around Paris is described as a degraded oceanic climate, the area around Lyon as a mixture of continental and mountain climates with a Mediterranean influence, and Toulouse as a south-western basin climate. It remains to be determined whether these different climates impact tick populations and their abundance, and whether this can be linked to the prevalence of equine piroplasmosis. In France, four tick species could be involved in the transmission of equine piroplasmosis agents: *Dermacentor reticulatus*, *D. marginatus*, *Hyalomma marginatum*, and *Rhipicephalus bursa* (ref Générale), and they have been found to be infesting horses in France (Chastagner et al., 2013; Vial et al., 2016; Grech-Angelini et al., 2016; Rocafort-Ferrer et al., 2022). *Hyalomma marginatum* and *Rhipicephalus bursa* are known to prefer southern climates (Estrada ?), and could therefore be more abundant in the southern part of France. However, data on their prevalence in horses in France is unknown.

A frequency of 38.3% of equine piroplasmosis carriers was assessed in the sampled population. The 4 geographic areas analyzed represent approximately one-third of the French equine population, with the Pays de la Loire and Auvergne-Rhône-Alpes being two of the four regions with the most abundant equine populations (IFCE, 2023). Our study lacks data on the French region with the largest equine population, Normandy, where prevalence is likely to be low because this breeding region is located in the northern part of France and a seroprevalence of less than 5% was determined using the CFT method (Nadal et al., 2022). However, one in three equid carrying piroplasmosis in France, with a gradient of about one in five in the north-west to one in two in the south-east of France, seem to be fairly realistic estimates of the prevalence of carriers of these pathogens.

Based on this molecular analysis of piroplasmosis carriers and questionnaire responses, we found a strong correlation between the frequency of carriers and cases of acute piroplasmosis reported in the four regions sampled. This correlation confirms the difference in circulation of the etiological agents of equine piroplasmosis between the four regions sampled. It also highlights the fact that owners are often not fully aware of the infectious status of their horses. The acquisition of an already infected equine or silent infections without clinical symptoms probably explain this difference between the frequency of carriage and the frequency of acute piroplasmosis. According to the questionnaires, diagnostics during acute piroplasmosis revealed *B. caballi* (17 cases) almost as often as *T. equi* (21 cases, including serological diagnostic) as the potential causal agent. In the diagnosis of acute piroplasmosis, *T. equi* can easily be mistaken as the causal agent of fever because of its lifelong persistence. When diagnosed during a fever, a *T. equi* carrier will be detected by serological and molecular means (this study), even if it is not the causal agent of the symptoms.

In France, the RESPE (Réseau d’Épidémio-surveillance en Pathologie Equine) provides data on equine piroplasmosis frequency through a combination of veterinary reports and molecular analysis of blood samples from horses with unexplained fever sent by sentinel veterinarians. On the samples collected and analyzed from 2013-2022 in France, *T. equi* was diagnosed in 34.6% (2,381 samples tested) and *B. caballi* in 13.9% (2,646 samples tested) of samples (Péju, 2023). The frequency of detection of *T. equi* was similar whether the sample consisted of asymptomatic or symptomatic horses (36.9% in our study and 34.6% in the RESPE analysis, respectively). However, the incidence of *B. caballi* differed, with 3.4% in asymptomatic horses compared to 13.9% in horses with a fever. This comparison suggests that the most symptomatic piroplasmosis cases likely stem from *B. caballi* infection, while the RESPE study population with *T. equi* detection likely represents carriers of a previous infection. This is further supported by the observation that equids with *T. equi* detection presented with milder clinical signs (fevers <39°C, weight loss, reduced performance, no anemia) compared to the severe signs (high fevers, anemia, weakness, anorexia, jaundice) associated with *B. caballi* infection (Péju, 2023). A quantitative analysis of the molecular detection could be particularly valuable in evaluating parasitemia levels in asymptomatic versus symptomatic horses.

The role of age as a risk factor for *T. equi* infection remains debatable, with studies supporting and contradicting this association (see discussion and references in Montes Cortès et al., 2017). These discrepancies may be linked to the endemicity level in each study area, and to the age class chosen in each study. In highly endemic regions, foals might acquire infections early through tick exposure or even by vertical transmission (Allsopp et al., 2007; Hermanns et al., 2024), leading to high prevalence as demonstrated in this study (more than one in four yearlings are *T. equi* carriers around Nantes, Toulouse or Lyon). Prevalence might reach a maximum at a young age, maximum that could correspond to the “environmental risk” (type of housing, tick species and abundance…). Conversely, in regions with low endemicity, the overall risk of infection (environmental risk) might be lower, making age a more significant factor. Older animals would then have a higher cumulative exposure to ticks and risk of transmission of pathogens over time. This explanation, of course, only applies to *T. equi* due to its lifelong carrier state in infected animals, unlike the different infection dynamics of *B. caballi*.

In this study, we sequenced partial 18S rRNA genes from most detected equine piroplasmosis agents, using the genotype references A to E for *T. equi* and A, B1 and B2 for *B. caballi* as described in the review by Tirosh-Levy et al., (2020). This analysis identified a predominant presence of the *T. equi* genotype E in France (98% of isolates, 193 sequences) across all studied regions. Only a few isolates (4) belonged to genotype A and were found scattered across different areas (near Lyon, Toulouse, and Paris). Data on *T. equi* sequences found in France is scarce. We often had to determine genotypes by performing bioinformatic analysis on GenBank sequence references, either deposited directly or referenced as containing identical sequences (Grech-Angelini et al., 2020; Fritz, 2010; Dahmana *et al.,* 2019; Joly Kukla et al., 2024; Bernard et al., 2024). Studies have shown genotype E to be present in all positive horse samples from the northern part of France (Fritz, 2010) and from Marseille on the Mediterranean coast (Dahmana *et al.,* 2019, *Theileria* sp. Europa var1). In Corsica, using standard conventional and real-time PCR methods, Dahmana et al. (2019) identified genotype A (46.1%, *T. equi* var2 and var3), genotype E (35.9%, *Theileria* sp. Europa var1), and genotype D (15.38%, *Theileria* sp. Africa var 4) in asymptomatic horses. In the South of France and in Corsica, using high-throughput BioMark microfluidic amplification, only genotype A was found in *H. marginatum* and *R. bursa* ticks collected from horses (Grech-Angelini et al., 2019; Joly Kukla et al., 2024; Bernard et al., 2024).

The genotype E is widespread in Eurasia (reviewed by Tirosh-Levy et al., 2020). Limited data from Eastern and Northern French neighboring countries highlights the presence of genotype E only in Ireland, Switzerland, Austria, Hungary and Czech Republic (Belkova et al., 2021; Liu et al., 2016; Dirks et al., 2021; Farkas et al., 2013; Hornok et al., 2020; Coultous et al., 2020). In contrast, southern European countries like Spain, Portugal, Italy, Croatia, Romania, and Greece typically show a mix of genotypes, including genotype E (Camino et al., 2020; Criado et al., 2006; Criado-Fornelio et al., 2004; Toma et al 2017; Veronesi et al., 2014; Chisu et al., 2023; Gallusova et al., 2014; Coultous et al., 2022; Fueher et al., 2020). This distribution of genotypes might be likely linked to the greater diversity of tick vectors present in southern Europe compared to northern regions.

In Spain, genotype E is well-represented, particularly in the north (Camino et al., 2020; Gimenez et al., 2009). Studies there have also suggested an association between genotype E and asymptomatic horses, while genotype A appears more frequent in symptomatic cases (Nagore et al., 2004; Criado-Fornelio et al., 2004; Criado et al., 2006). As our study focused on asymptomatic horses in order to evaluate the equine piroplasmosis reservoir, it should be complemented by a similar study on symptomatic horses in order to assess any potential bias this might introduce into our genetic diversity findings. Two of the four horses carrying the *T. equi* A genotype originated in Spain. They likely acquired this variant there and remained carriers.

All 19 *B. caballi* sequences obtained in this study belonged to genotype A, which is the most prevalent genotype worldwide (Tirosh-Levy et al., 2020). However, due to the low number of *B. caballi* carriers in our study, other genotypes may still exist in France. Genotypes B1 and B2 have been described in *H. marginatum* and *R. annulatus* ticks in Italy (Toma et al., 2017) and B1 has been found in horses in Spain (Criado-Fornelio et al., 2004; Nagore et al., 2004; Criado et al., 2006; Camino et al., 2020). To gain a more comprehensive understanding of *B. caballi* genetic diversity in France, we would need to increase sampling of either symptomatic horses or ticks.

## Conclusion

Equine piroplasmosis is endemic in France with a carrier frequency of 38.3%, frequency of *T. equi* carriers higher in the southern parts of the country, and a low frequency of *B. caballi* carriers. No significant effect of horse age or gender on these frequencies could be demonstrated. The genetic diversity of isolates in France is rather low, with a great majority of *T. equi* genotype E and *B. caballi* genotype A.

## CRediT authorship contribution statement

**Maggy Jouglin**: Project Administration – Formal analysis - Investigation - Writing: Review & Editing. **Claire Bonsergent**: Investigation - Formal analysis - Writing: Review & Editing. **Nathalie de la Cotte**: Investigation - Writing: Review & Editing. **Mickaël Mège**: Investigation - Formal Analysis – Visualization - Writing: Review & Editing. **Céline Bizon**: Investigation - Resources - Writing: Review & Editing. **Anne Couroucé**: Resources - Project Administration - Writing: Review & Editing. **Elodie Lallemand**: Investigation - Project Administration - Resources - Writing: Review & Editing. **Louise Lemonnier**: Investigation - Resources - Writing: Review & Editing. **Agnès Leblond**: Investigation - Project Administration - Resources - Writing: Review & Editing. **Aurélia Leroux:** Investigation - Resources - Writing: Review & Editing. **Ilaria Marano**: Investigation - Resources - Writing: Review & Editing. **Alexandre Muzard**: Resources - Formal Analysis - Writing: Review & Editing. **Emilie Quéré**: Investigation - Project Administration - Resources - Writing: Review & Editing. **Marion Toussaint**: Investigation - Resources - Project Administration - Writing: Review & Editing. **Albert Agoulon:** Formal Analysis - Funding Acquisition – Visualization - Writing: Review & Editing. **Laurence Malandrin:** Conceptualization - Project Administration - Funding Acquisition – Supervision – Visualization - Writing: Original Draft Preparation, Review & Editing.

## Declaration of generative AI and AI-assisted technologies in the writing process

During the preparation of this work, LM used Gemini in order to improve the English language. After using this tool, the authors reviewed and edited the content as needed and take full responsibility for the content of the publication.

## Declaration of competing interest

The authors declare that they have no competing interests.

## Data availability

The datasets used and/or analyzed during the current study are available from L. Malandrin on reasonable request.

## Acknowledgments

We wish to thank Aymeric Bohec, Olivia Guibert, Alexandra Prévôt, and Marthe Thirouin for their unvaluable help in the blood collection

## Supplementary materials

Supplementary tables 1 to 6 and supplementary figures 1 to 3 are available at ….

## Funding

The study was funded by IFCE (Institut Français du Cheval et de l’Equitation [French Institute for Horses and Riding]), Fonds Eperon and Institut Carnot France Futur Elevage.

## Notes

### Competing Interest Statement

The authors have declared no competing interest.

