## Supplementary tables for "Equine piroplasmosis in different geographical areas in France: prevalence heterogeneity of asymptomatic carriers and low genetic diversity of *Theileria equi* and *Babesia caballi*"

Supplementary table 1. Sampled population and prevalence of asymptomatic carriers of piroplasmosis, *T. equi* and *B. caballi* in equids sampled in the 4 veterinary schools between 2019 and 2023.

| Veterinary School | Year **Period** | Number of sampled equids | Carrier frequency % [95% CI] (number) of | | | |
|---|---|---|---|---|---|---|
| | | | Piroplasmosis carriers | *T. equi* carriers | *B. caballi* carriers | Co-infected carriers |
| Paris | 2020 | 39 | 15.4 [6.4-31.2] (6) | 10.3 [3.3-25.2] (4) | 5.1 [0.9-18.6] (2) | 0 [0.0-11.2] (0) |
| | 2021 | 53 | 24.5 [14.2-38.6] (13) | 24.5 5 [14.2-38.6] (13) | 0 [0.0-8.4] (0) | 0 [0.0-8.4] (0) |
| | 2022 | 15 | 6.7 [0.3-34.0] (1) | 6.7 [0.3-34.0] (1) | 0 [0.0-25.3] (0) | 0 [0.0-25.3] (0) |
| | **2020-2022** | **107** | **18.7 [12.1-27.6] (20)** | **16.8 [10.5-25.6] (18)** | **1.9 [0.3-7.2] (2)** | **0 [0.0-4.3] (0)** |
| Nantes | 2019 | 71 | 26.8 [17.3-38.8] (19) | 26.8 [17.3-38.8] (19) | 0 [0.0-6.4] (0) | 0 [0.0-6.4] (0) |
| | 2020 | 62 | 40.3 [28.3-53.5] (25) | 40.3 [28.3-53.5] (25) | 0 [0.0-7.3] (0) | 0 [0.0-7.3] (0) |
| | 2021 | 32 | 31.3 [16.7-50.1] (10) | 31.3 [16.7-50.1] (10) | 0 [0.0-13.3] (0) | 0 [0.0-13.3] (0) |
| | 2022 | 0 | 0 | 0 | 0 | 0 |
| | 2023 | 4 | 0 [0.0-60.4] (0) | 0 [0.0-60.4] (0) | 0 [0.0-60.4] (0) | 0 [0.0-60.4] (0) |
| | **2019-2023** | **169** | **32.0 [25.1-39.6] (54)** | **32.0 [25.1-39.6] (54)** | **0 [0.0-2.8] (0)** | **0 [0.0-2.8] (0)** |
| Toulouse | 2020 | 5 | 40.0 [7.3-83.0] (2) | 40.0 [7.3-83.0] (2) | 0 [0.0-53.7] (0) | 0 [0.0-53.7] (0) |
| | 2021 | 52 | 48.1 [34.2-62.2] (25) | 46.2 [32.5-60.4] (24) | 5.8 [1.5-16.9] (3) | 3.8 [0.7-14.3] (2) |
| | 2022 | 78 | 37.2 [26.7-48.9] (29) | 37.2 [26.7-48.9] (29) | 5.1 [1.7-13.3] (4) | 5.1 [1.7-13.3] (4) |
| | **2020-2022** | **135** | **41.5 [33.2-50.3] (56)** | **40.7 [32.5-49.5] (55)** | **5.2 [2.3-10.8] (7)** | **4.4 [1.8-9.8] (6)** |
| Lyon | 2020 | 18 | 44.4 [22.4-68.6] (8) | 44.4 [22.4-68.6] (8) | 0 [0.0-21.9] (0) | 0 [0.0-21.9] (0) |
| | 2021 | 87 | 62.1 [51.0-72.1] (54) | 57.5 [46.4-67.9] (50) | 8.0 [3.6-16.4] (7) | 3.4 [0.9-10.5] (3) |
| | 2022 | 50 | 50.0 [36.6-63.4] (25) | 48.0 [33.9-62.4] (24) | 6.0 [1.6-17.5] (3) | 4.0 [0.7-14.9] (2) |
| | **2020-2022** | **155** | **56.1 [47.9-64.0] (87)** | **52.9 [44.8-60.9] (82)** | **6.5 [3.3-11.9] (10)** | **3.2 [1.2-7.8] (5)** |
| **Studied population** | **2019-2023** | **566** | **38.3 [34.3-42.5] (217)** | **36.9 [33.0-41.1] (209)** | **3.4 [2.1-5.3] (19)** | **1.9 [1.0-3.6] (11)** |

Supplementary Table 2: Number of tested and piroplasmosis carrier horses according to the departments and regions and calculated prevalence by region (data used to generate the Figure 2 in the manuscript and for the statistical analysis). The colors used in the table correspond to the colors used in the figure 2. The regions in the north of France are in the left table and those in the south of France in the right table.

| County number | Nb of tested equids | Nb of piroplasmosis carrier equids | Prevalence (region scale) |
|---|---|---|---|
| 02 | 1 | 0 | 2/13 (15.4%) |
| 59 | 2 | 0 | |
| 60 | 9 | 1 | |
| 62 | 1 | 1 | |
| **80** | 0 | 0 | |
| 14 | 2 | 0 | 0/8 |
| 27 | 4 | 0 | |
| 76 | 2 | 0 | |
| **50, 61** | 0 | 0 | |
| 75 | 5 | 3 | 12/68 (17.65%) |
| 77 | 10 | 2 | |
| 78 | 26 | 4 | |
| 91 | 13 | 1 | |
| 92 | 2 | 0 | |
| 94 | 4 | 1 | |
| 95 | 8 | 1 | |
| **93** | 0 | 0 | |
| 54 | 2 | 0 | 0/2 |
| **08, 10, 51, 52, 55, 57, 67, 68, 88** | 0 | 0 | |
| 22 | 2 | 0 | 7/29 (24.1%) |
| 29 | 2 | 0 | |
| 35 | 12 | 2 | |
| 56 | 13 | 5 | |
| 44 | 100 | 31 | 39/120 (32.5%) |
| 49 | 10 | 5 | |
| 85 | 10 | 3 | |
| **53, 72** | 0 | 0 | |
| 18 | 1 | 1 | 11/26 (42.31%) |
| 28 | 2 | 2 | |
| 37 | 3 | 1 | |
| 41 | 15 | 6 | |
| 45 | 5 | 1 | |
| 21 | 3 | 2 | 14/17 (82.35%) |
| 39 | 3 | 3 | |
| 70 | 1 | 1 | |
| 71 | 7 | 6 | |
| 89 | 3 | 2 | |

| County number | Nb of tested equids | Nb of piroplasmosis carrier equids | Prevalence (region scale) |
|---|---|---|---|
| 16 | 1 | 0 | 6/22 (27.27%) |
| 24 | 4 | 1 | |
| 33 | 5 | 2 | |
| 40 | 1 | 0 | |
| 47 | 4 | 1 | |
| 64 | 1 | 0 | |
| 79 | 6 | 2 | |
| **17, 19, 23, 86, 87** | 0 | 0 | |
| 01 | 10 | 6 | 68/131 (51.91%) |
| 07 | 1 | 0 | |
| 26 | 5 | 3 | |
| 38 | 23 | 12 | |
| 42 | 8 | 6 | |
| 43 | 3 | 2 | |
| 63 | 1 | 1 | |
| 69 | 74 | 36 | |
| 73 | 2 | 1 | |
| 74 | 4 | 1 | |
| **03, 15** | 0 | 0 | |
| 09 | 4 | 3 | 52/116 (44.83%) |
| 11 | 3 | 1 | |
| 12 | 3 | 1 | |
| 30 | 3 | 1 | |
| 31 | 47 | 22 | |
| 32 | 10 | 5 | |
| 46 | 8 | 5 | |
| 48 | 1 | 1 | |
| 65 | 10 | 5 | |
| 66 | 4 | 1 | |
| 81 | 11 | 4 | |
| 82 | 12 | 3 | |
| **34** | 0 | 0 | |
| 83 | 1 | 0 | 0/1 |
| **04 , 05, 06, 13, 84** | 0 | 0 | |
| 02A, 02B | 0 | 0 | 0 |

☐ Hauts-de-France ☐ Normandy ☐ Ile-de-France ☐ Grand-Est
☐ Brittany ☐ Pays de la Loire ☐ Centre-Val de Loire ☐ Burgundy-Franche-Comté
☐ New-Aquitaine ☐ Auvergne-Rhône-Alpes
☐ Occitanie ☐ Provence-Alpes-Côte d'Azur ☐ Corsica

Supplementary table 3. Statistical analysis of equine piroplasmosis carrier frequencies between the French regions. p-values of 2 by 2 frequency comparisons of carriers using the Fisher's exact test are indicated. The significance threshold was set at 0.0014 after the Bonferroni correction and the significant differences are highlighted in bold.

| | Pays de la Loire | Hauts-de-France | Ile-de-France | Centre-Val de Loire | Burgundy-Franche-Comté | New-Aquitaine | Auvergne-Rhône-Alpes | Occitanie |
|---|---|---|---|---|---|---|---|---|
| Brittany (7/29) 24.1% | 0.5028 | 0.6953 | 0.577 | 0.2494 | **0.0002** | 1 | 0.0075 | 0.0567 |
| Pays de la Loire (39/120) 32.5% | | 0.3429 | 0.0398 | 0.3671 | **0.0001** | 0.8041 | 0.0022 | 0.0613 |
| Hauts-de-France (2/13) 15.4% | | | 1 | 0.1514 | **0.0006** | 0.68 | 0.0175 | 0.0719 |
| Ile-de-France (12/68) 17.6% | | | | 0.0174 | **1.025e-06** | 0.3639 | 1.978e-06 | **0.0002** |
| Centre-Val de Loire (11/26) 42.3% | | | | | 0.0125 | 0.3681 | 0.3982 | 1 |
| Burgundy-Franche-Comté (14/17) 82.4% | | | | | | **0.0011** | 0.0199 | 0.0042 |
| New-Aquitaine (6/22) 27.3% | | | | | | | 0.0388 | 0.1598 |
| Auvergne-Rhône-Alpes (68/131) 51.9% | | | | | | | | 0.3079 |
| Occitanie (52/116) 44.8% | | | | | | | | |

Supplementary table 4: *T. equi* genotype E sequence details and similarities with Genbank reference sequences typed as genotype E (AY534882 and KF559357) and genotype A (KJ573370).

| Isolate reference | County number | Sequence length | Genbank accession number | AY534882 E - Spain | KF559357 E - China | KJ573370 A- Brazil |
|---|---|---|---|---|---|---|
| CV19-07 | 44 | 1115 | | 99.37 | 99.64 | 96.42 |
| CV19-08 | 44 | 818 | | 99.63 | 99.63 | 94.88 |
| CV19-16 | 44 | 1267 | | 99.76 | 100.00 | 96.44 |
| CV19-41 | 44 | 475 | | 99.58 | 99.58 | 93.08 |
| CV19-42 | 44 | 1066 | | 99.72 | 100.00 | 96.35 |
| CV19-44 | 44 | 607 | | 99.01 | 99.34 | 94.25 |
| CV19-45 | 44 | 1371 | PQ044745 | 99.78 | 100.00 | 96.91 |
| CV19-46 | 56 | 718 | | 99.72 | 99.72 | 96.13 |
| CV19-48 | 79 | 1390 | PQ044746 | 99.71 | 99.93 | 96.93 |
| CV19-49 | 37 | 1394 | PQ044747 | 99.79 | 100.00 | 96.96 |
| CV19-50 | 44 | 1398 | | 99.79 | 100.00 | 96.95 |
| CV19-51 | 44 | 1389 | | 99.78 | 100.00 | 96.93 |
| CV19-52 | 44 | 1150 | | 99.83 | 100.00 | 96.43 |
| CV19-56 | 49 | 1382 | | 99.71 | 99.93 | 96.90 |
| CV19-60 | 85 | 1406 | | 99.72 | 99.93 | 96.96 |
| CV19-61 | 44 | 409 | | 99.27 | 99.27 | 94.54 |
| CV19-62 | 49 | 1405 | PQ044748 | 99.79 | 100.00 | 96.96 |
| CV19-73 | 44 | 1404 | | 99.79 | 100.00 | 96.95 |
| CV19-75 | 44 | 551 | | 99.46 | 99.46 | 97.61 |
| CV20-02 | 85 | 1399 | PQ044749 | 99.79 | 100.00 | 96.94 |
| CV20-03 | 56 | 1407 | PQ044750 | 99.79 | 100.00 | 96.95 |
| CV20-10 | 44 | 1412 | | 99.79 | 100.00 | 96.96 |
| CV20-15 | 85 | 1380 | | 99.49 | 99.71 | 96.91 |
| CV20-18 | 49 | 1397 | | 99.79 | 100.00 | 96.93 |
| CV20-21 | 44 | 720 | | 99.58 | 99.72 | 95.42 |
| CV20-25 | 56 | 1372 | | 99.78 | 100.00 | 96.91 |
| CV20-31 | 41 | 1389 | | 99.71 | 99.93 | 96.93 |
| CV20-32 | 41 | 1393 | | 99.79 | 100.00 | 96.96 |
| CV20-33 | 41 | 777 | | 99.61 | 100.00 | 95.89 |
| CV20-34 | 41 | 1363 | | 99.78 | 100.00 | 96.91 |
| CV20-38 | 41 | 1400 | PQ044751 | 99.71 | 99.93 | 96.94 |
| CV20-51 | 44 | 1394 | | 99.79 | 100.00 | 96.93 |
| CV20-53 | 44 | 794 | | 99.62 | 100.00 | 96.11 |
| CV20-56 | 35 | 820 | | 99.64 | 100.00 | 96.11 |
| CV20-61 | 79 | 787 | | 99.62 | 100.00 | 95.94 |
| CV20-62 | 49 | 831 | | 99.64 | 100.00 | 96.15 |
| CV20-63 | 44 | 818 | | 99.63 | 100.00 | 96.10 |
| CV20-64 | 44 | 803 | | 99.63 | 100.00 | 96.02 |
| CV20-65 | 35 | 983 | PQ044752 | 99.70 | 100.00 | 96.65 |
| CV20-66 | 44 | 842 | | 99.64 | 100.00 | 96.21 |
| CV20-68 | 56 | 930 | | 99.68 | 100.00 | 96.57 |
| CV20-71 | 44 | 833 | | 99.64 | 100.00 | 96.25 |
| CV20-73 | 44 | 932 | | 99.57 | 99.89 | 96.17 |
| CV20-75 | 56 | 825 | | 99.64 | 100.00 | 96.47 |
| CV21-05 | 44 | 1380 | | 99.71 | 99.93 | 96.86 |
| CV21-08 | 49 | 1380 | | 99.71 | 99.93 | 96.91 |
| CV21-15 | 44 | 1378 | | 99.57 | 99.78 | 96.91 |
| CV21-17 | 44 | 890 | | 100.00 | 100.00 | 98.51 |
| CV21-21 | 44 | 819 | | 100.00 | 100.00 | 98.52 |
| CV21-26 | 44 | 1394 | | 99.78 | 100.00 | 97.03 |
| CV21-27 | 44 | 1402 | | 99.71 | 99.93 | 97.05 |
| CV21-29 | 44 | 829 | | 99.64 | 100.00 | 96.15 |
| CVA20-13 | 45 | 920 | PQ044753 | 99.57 | 99.89 | 96.42 |
| CVA20-35 | 95 | 868 | | 99.66 | 100.00 | 96.32 |
| CVA20-40 | 37 | 813 | | 99.63 | 100.00 | 96.07 |
| CVA21-02 | 75 | 1381 | | 99.78 | 100.00 | 96.93 |
| CVA21-07 | 78 | 1390 | | 99.71 | 99.93 | 96.93 |
| CVA21-08 | 78 | 1389 | PQ044754 | 99.78 | 100.00 | 96.93 |

| | | | | | | |
|---|---|---|---|---|---|---|
| CVA21-09 | 89 | 1388 | PQ044755 | 99.78 | 100.00 | 96.93 |
| CVA21-11 | 77 | 1378 | PQ044756 | 99.78 | 100.00 | 96.93 |
| CVA21-12 | 77 | 1381 | | 99.78 | 100.00 | 96.93 |
| CVA21-16 | 78 | 890 | | 99.66 | 100.00 | 96.41 |
| CVA21-20 | 60 | 1390 | PQ044757 | 99.71 | 99.93 | 96.86 |
| CVA21-31 | 94 | 1395 | | 99.79 | 100.00 | 97.04 |
| CVA21-39 | 78 | 851 | | 99.65 | 100.00 | 96.25 |
| CVA21-45 | 75 | 1375 | | 99.78 | 100.00 | 96.91 |
| CVA21-50 | 62 | 800 | PQ044758 | 99.63 | 100.00 | 96.01 |
| CVA21-53 | 94 | 845 | | 99.65 | 100.00 | 96.22 |
| CVA21-74 | 41 | 696 | | 100.00 | 100.00 | 98.54 |
| CVT20-01 | 32 | 816 | | 99.63 | 100.00 | 96.09 |
| CVT20-12 | 31 | 830 | | 99.64 | 100.00 | 96.15 |
| CVT21-02 | 46 | 847 | | 99.65 | 100.00 | 96.23 |
| CVT21-03 | 81 | 1389 | PQ044759 | 99.78 | 100.00 | 96.93 |
| CVT21-06 | 31 | 1007 | | 99.90 | 99.90 | 95.76 |
| CVT21-07 | 12 | 1026 | PQ044760 | 99.90 | 99.90 | 95.83 |
| CVT21-08 | 31 | 842 | | 100.00 | 100.00 | 98.55 |
| CVT21-09 | 33 | 974 | PQ044761 | 99.90 | 99.90 | 96.85 |
| CVT21-15 | 65 | 854 | | 99.65 | 100.00 | 96.26 |
| CVT21-16 | 65 | 1395 | | 99.79 | 100.00 | 97.04 |
| CVT21-19 | 24 | 853 | PQ044762 | 99.65 | 100.00 | 96.26 |
| CVT21-28 | 82 | 828 | | 99.64 | 100.00 | 96.15 |
| CVT21-29 | 31 | 831 | | 99.64 | 100.00 | 96.16 |
| CVT21-32 | nd | 1021 | | 99.90 | 99.90 | 95.84 |
| CVT21-33 | 65 | 917 | PQ044763 | 99.78 | 99.78 | 98.23 |
| CVT21-35 | 31 | 908 | | 99.67 | 100.00 | 96.48 |
| CVT21-36 | 31 | 1388 | PQ044764 | 99.78 | 100.00 | 96.93 |
| CVT21-38 | 31 | 971 | | 100.00 | 100.00 | 96.95 |
| CVT21-39 | 31 | 934 | | 99.68 | 100.00 | 96.58 |
| CVT21-40 | 31 | 943 | | 99.68 | 100.00 | 96.61 |
| CVT21-42 | 82 | 835 | PQ044765 | 99.64 | 100.00 | 96.18 |
| CVT21-47 | 32 | 1073 | PQ044766 | 99.72 | 100.00 | 96.83 |
| CVT21-48 | 31 | 951 | | 99.69 | 100.00 | 96.64 |
| CVT21-50 | 30 | 931 | PQ044767 | 99.68 | 100.00 | 96.57 |
| CVT21-55 | 32 | 919 | | 99.67 | 100.00 | 96.53 |
| CVT21-58 | 31 | 961 | | 99.27 | 99.27 | 96.31 |
| CVT21-59 | 47 | 1101 | PQ044768 | 99.91 | 99.91 | 96.09 |
| CVT21-62 | 46 | 964 | PQ044769 | 99.79 | 99.79 | 96.95 |
| CVT21-77 | 66 | 1001 | PQ044770 | 99.30 | 99.30 | 95.36 |
| CVT21-78 | 09 | 880 | | 99.66 | 100.00 | 96.37 |
| CVT22-02 | 31 | 1371 | | 99.71 | 99.93 | 96.91 |
| CVT22-05 | 46 | 753 | | 99.60 | 100.00 | 95.76 |
| CVT22-06 | 82 | 450 | | 99.34 | 100.00 | 96.33 |
| CVT22-15 | 31 | 872 | | 100.00 | 100.00 | 98.48 |
| CVT22-16 | 31 | 558 | | 99.28 | 99.82 | 94.46 |
| CVT22-19 | 81 | 918 | | 99.67 | 100.00 | 96.52 |
| CVT22-21 | 65 | 884 | | 99.66 | 100.00 | 96.39 |
| CVT22-22 | 46 | 552 | | 99.46 | 100.00 | 94.59 |
| CVT22-23 | 31 | 816 | | 99.63 | 100.00 | 96.09 |
| CVT22-24 | 31 | 749 | | 100.00 | 100.00 | 98.52 |
| CVT22-29 | 31 | 459 | | 99.35 | 100.00 | 93.93 |
| CVT22-33 | 31 | 489 | | 99.56 | 100.00 | 95.36 |
| CVT22-34 | 81 | 687 | | 99.39 | 100.00 | 94.09 |
| CVT22-35 | 33 | 935 | | 99.68 | 100.00 | 96.59 |
| CVT22-37 | 65 | 499 | | 99.80 | 99.80 | 98.14 |
| CVT22-38 | 31 | 807 | | 100.00 | 100.00 | 98.51 |
| CVT22-39 | 46 | 783 | | 99.62 | 100.00 | 96.05 |
| CVT22-40 | 31 | 703 | | 99.86 | 99.86 | 98.52 |
| CVT22-41 | 09 | 1357 | PQ044771 | 99.71 | 99.93 | 96.88 |
| CVT22-49 | 32 | 812 | | 99.63 | 100.00 | 96.19 |
| T19-1347 | 81 | 1352 | | 99.78 | 100.00 | 96.96 |
| CVL20-01 | 01 | 824 | | 99.64 | 100.00 | 96.25 |
| CVL20-04 | 71 | 906 | | 99.67 | 100.00 | 96.59 |

| | | | | | | |
|---|---|---|---|---|---|---|
| CVL20-08 | 38 | 960 | | 99.48 | 99.79 | 96.57 |
| CVL20-10 | 38 | 831 | | 99.64 | 100.00 | 96.27 |
| CVL20-14 | 01 | 230 | | 99.57 | 99.57 | 87.39 |
| CVL20-15 | 71 | 1395 | | 99.64 | 99.86 | 96.89 |
| CVL20-16 | 38 | 908 | PQ044772 | 99.67 | 100.00 | 96.59 |
| CVL20-18 | 01 | 866 | | 99.54 | 99.89 | 96.31 |
| CVL20-21 | nd | 830 | | 99.64 | 100.00 | 96.27 |
| CVL20-22 | nd | 799 | | 99.63 | 100.00 | 96.13 |
| CVL21-01 | 71 | 831 | | 99.64 | 100.00 | 96.27 |
| CVL21-02b | 69 | 1040 | | 100.00 | 100.00 | 96.16 |
| CVL21-02c | 69 | 1371 | | 99.35 | 99.56 | 96.57 |
| CVL21-03 | 69 | 1388 | PQ044773 | 99.78 | 100.00 | 96.98 |
| CVL21-05 | 69 | 1392 | | 99.71 | 99.93 | 97.00 |
| CVL21-07 | 01 | 1361 | | 99.78 | 100.00 | 96.96 |
| CVL21-08 | 42 | 833 | | 99.64 | 100.00 | 96.27 |
| CVL21-09 | 69 | 969 | | 99.28 | 99.28 | 97.70 |
| CVL21-13 | 69 | 1381 | | 99.78 | 100.00 | 97.01 |
| CVL21-16 | 42 | 1393 | | 99.79 | 100.00 | 97.00 |
| CVL21-17 | 42 | 1398 | | 99.79 | 100.00 | 97.01 |
| CVL21-21 | 26 | 1385 | | 99.78 | 100.00 | 97.01 |
| CVL21-22 | 69 | 1410 | | 99.72 | 99.93 | 97.03 |
| CVL21-23 | 69 | 882 | | 99.43 | 99.43 | 97.85 |
| CVL21-24 | nd | 1386 | | 99.78 | 100.00 | 97.01 |
| CVL21-25 | 69 | 1409 | | 99.72 | 99.93 | 97.03 |
| CVL21-26 | 69 | 1408 | | 99.72 | 99.93 | 97.03 |
| CVL21-27 | 69 | 1045 | | 100.00 | 100.00 | 96.10 |
| CVL21-31 | 39 | 990 | PQ044774 | 99.39 | 99.39 | 96.54 |
| CVL21-32 | 69 | 962 | | 99.27 | 99.27 | 96.53 |
| CVL21-33 | 69 | 1026 | | 100.00 | 100.00 | 96.02 |
| CVL21-34 | 43 | 1044 | PQ044775 | 100.00 | 100.00 | 96.12 |
| CVL21-35 | 69 | 921 | | 99.89 | 99.89 | 96.84 |
| CVL21-36 | 43 | 916 | | 100.00 | 100.00 | 98.45 |
| CVL21-37 | nd | 980 | | 99.90 | 99.90 | 96.95 |
| CVL21-38 | 69 | 1017 | | 99.80 | 99.80 | 95.83 |
| CVL21-39 | 42 | 1045 | | 99.90 | 99.90 | 96.05 |
| CVL21-40 | 38 | 889 | | 99.78 | 99.78 | 98.51 |
| CVL21-41 | 69 | 944 | | 99.89 | 99.89 | 98.25 |
| CVL21-42 | 38 | 874 | | 100.00 | 100.00 | 98.49 |
| CVL21-43 | 69 | 969 | | 98.97 | 98.97 | 97.37 |
| CVL21-44 | 42 | 1035 | | 99.42 | 99.42 | 95.55 |
| CVL21-53 | 38 | 1374 | | 99.78 | 100.00 | 96.99 |
| CVL21-54 | 38 | 1394 | | 99.71 | 99.93 | 96.96 |
| CVL21-56 | 71 | 923 | | 99.68 | 100.00 | 96.65 |
| CVL21-57 | 69 | 1073 | | 100.00 | 100.00 | 96.23 |
| CVL21-58 | 26 | 1367 | PQ044776 | 99.78 | 100.00 | 96.98 |
| CVL21-59 | 71 | 1381 | PQ044777 | 99.78 | 100.00 | 97.01 |
| CVL21-63 | 42 | 1381 | PQ044778 | 99.78 | 100.00 | 97.01 |
| CVL21-67 | 69 | 512 | | 99.41 | 100.00 | 94.54 |
| CVL21-68 | 70 | 829 | PQ044779 | 99.64 | 100.00 | 96.27 |
| CVL21-83 | 69 | 919 | | 99.67 | 100.00 | 96.63 |
| CVL21-84 | 69 | 923 | | 99.68 | 100.00 | 96.65 |
| CVL21-85 | 63 | 923 | PQ044780 | 99.68 | 100.00 | 96.65 |
| CVL21-86 | 69 | 874 | | 99.54 | 99.89 | 96.35 |
| CVL22-01 | 69 | 495 | | 100.00 | 100.00 | 97.92 |
| CVL22-08 | 38 | 728 | | 99.59 | 100.00 | 95.75 |
| CVL22-09 | 38 | 749 | | 99.60 | 100.00 | 95.87 |
| CVL22-10 | 69 | 669 | | 99.40 | 99.85 | 95.38 |
| CVL22-11 | 69 | 916 | | 99.67 | 100.00 | 96.62 |
| CVL22-12 | 69 | 732 | | 99.32 | 99.73 | 95.50 |
| CVL22-17 | 48 | 1393 | | 99.78 | 100.00 | 97.03 |
| CVL22-18 | 71 | 912 | | 99.67 | 100.00 | 96.61 |
| CVL22-19 | 69 | 559 | | 99.46 | 100.00 | 94.83 |
| CVL22-20 | nd | 456 | | 99.34 | 100.00 | 96.24 |
| CVL22-23 | 69 | 689 | | 99.57 | 100.00 | 95.51 |

| | | | | | | |
|---|---|---|---|---|---|---|
| CVL22-27 | 01 | 628 | | 99.52 | 100.00 | 95.24 |
| CVL22-30 | 38 | 792 | | 99.34 | 99.74 | 95.64 |
| CVL22-31 | 69 | 550 | | 99.46 | 100.00 | 94.75 |
| CVL22-33 | 38 | 815 | | 99.63 | 100.00 | 96.21 |
| CVL22-34 | 74 | 688 | | 100.00 | 100.00 | 98.53 |
| CVL22-36 | 21 | 916 | PQ044781 | 99.67 | 100.00 | 96.62 |
| CVL22-37 | 21 | 909 | | 100.00 | 100.00 | 98.45 |
| CVL22-40 | 69 | 809 | | 99.63 | 100.00 | 96.18 |
| CVL22-48 | 38 | 1402 | | 99.71 | 99.93 | 96.98 |
| CVL22-51 | 69 | 1392 | | 99.78 | 100.00 | 97.03 |
| CVL22-52 | 01 | 810 | PQ044782 | 99.63 | 100.00 | 96.18 |

Supplementary table 5: *T. equi* genotype A sequence details and similarities with Genbank genotype A reference sequences.

| | County number | Sequence length | Genbank accession number | AY150062 A Spain | KJ573370 A Brazil | KX227625 A Israel |
|---|---|---|---|---|---|---|
| CVA20-26 | 89 | 1382 | PQ044783 | 99.64 | 99.93 | 99.86 |
| CVT21-14 | 32 | 998 | PQ044784 | 99.00 | 99.29 | 99.10 |
| CVT21-49 | 31 | 819 | PQ044785 | 99.76 | 99.88 | 100 |
| CVL21-18 | nd | 1393 | PQ044786 | 98.99 | 99.27 | 99.00 |

Supplementary table 6: *B. caballi* sequence details and similarities with Genbank reference sequences genotyped as genotype A (KY952238 and AY534883) and genotype B1 (EU642513).

| | County number | Sequence length | Genbank accession number | KY952238 A- Brazil | AY534883 A- Spain | EU642513 B1 South Africa |
|---|---|---|---|---|---|---|
| CVA20-01 | 75 | 824 | | 100 | 99.76 | 97.21 |
| CVA20-25 | 18 | 958 | PQ044787 | 100 | 99.79 | 97.97 |
| CVL21-06 | 69 | 815 | | 100 | 99.88 | 97.18 |
| CVL21-43 | 69 | 757 | | 100 | 100 | 95.95 |
| CVL21-44 | 42 | 779 | | 100 | 99.87 | 97.04 |
| CVL21-55 | 69 | 817 | PQ044788 | 100 | 99.76 | 97.18 |
| CVL21-56 | 71 | 765 | PQ044789 | 100 | 99.87 | 96.99 |
| CVL21-62 | 69 | 825 | | 100 | 99.76 | 97.21 |
| CVL21-64 | 73 | 815 | PQ044790 | 100 | 99.88 | 97.18 |
| CVL22-17 | 42 | 924 | PQ044791 | 100 | 99.78 | 97.95 |
| CVL22-47 | 69 | 765 | | 100 | 99.87 | 96.60 |
| CVL22-48 | 38 | 817 | | 100 | 99.76 | 97.31 |
| CVT21-14 | 32 | 822 | | 100 | 99.76 | 97.20 |
| CVT21-43 | 09 | 764 | PQ044792 | 99.21 | 99.08 | 97.12 |
| CVT21-48 | 31 | 765 | PQ044793 | 99.74 | 99.61 | 96.86 |
| CVT21-55 | 32 | 866 | PQ044794 | 100 | 99.77 | 98.15 |
| CVT21-62 | 46 | 927 | PQ044795 | 99.89 | 99.68 | 96.44 |
| CVT22-05 | 46 | 687 | | 100 | 99.71 | 98.46 |
| CVT22-16 | 31 | 651 | | 100 | 99.69 | 98.31 |
