## Supplementary figures for "Equine piroplasmosis in different geographical areas in France: prevalence heterogeneity of asymptomatic carriers and low genetic diversity of *Theileria equi* and *Babesia caballi*"

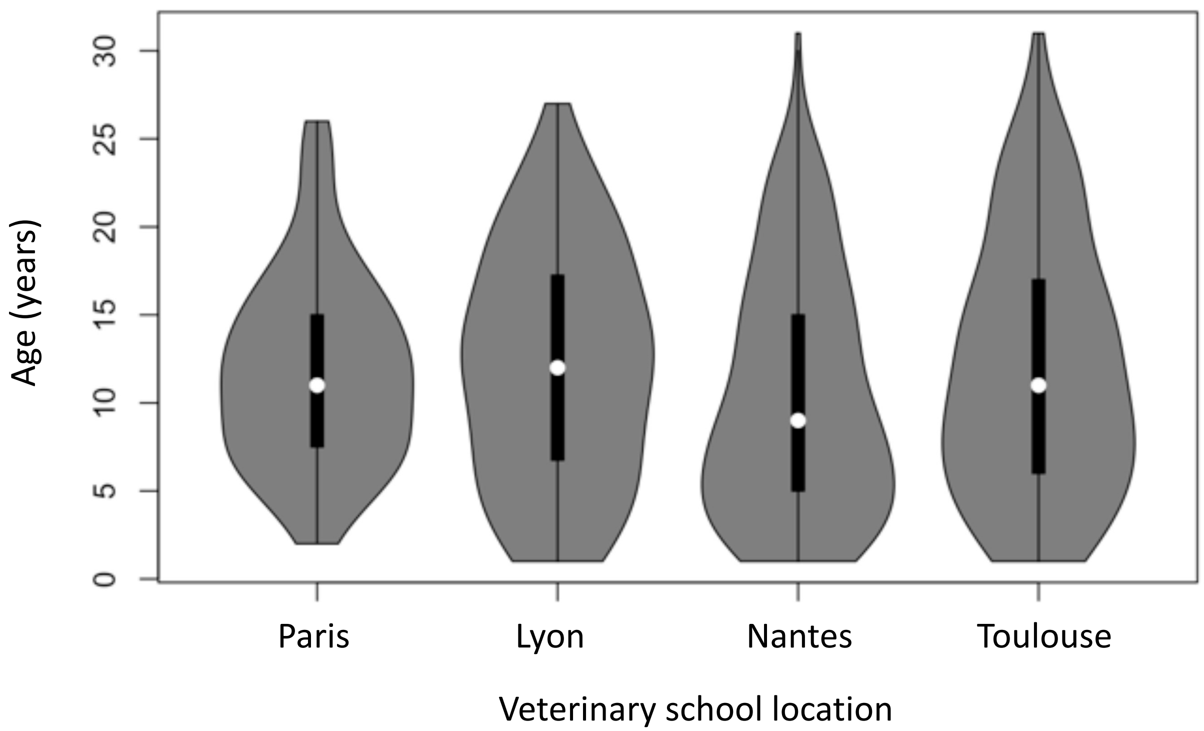


Supplementary Fig. 1. Violin diagram of the horse population age in the 4 sampled areas.

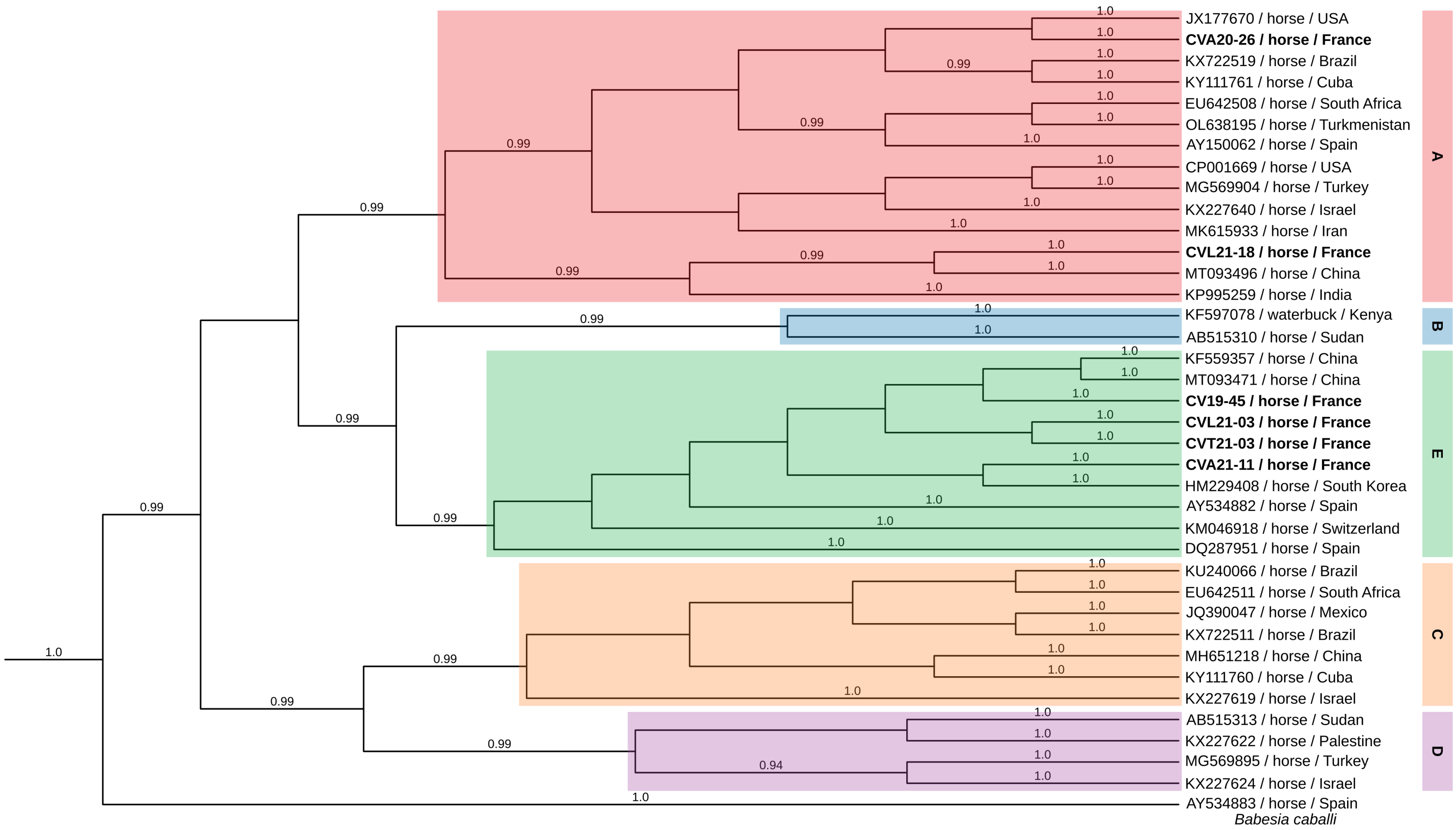


Supplementary Fig. 2. Phylogenetic tree of the 18S rRNA gene partial sequences of *Theileria equi* constructed by Bayesian inference and represented as a cladogram. The tree comprises 38 sequences for which 1295 positions were analyzed. Sequences from this study are in bold and the five genotypes (A, B, C, D and E) have been highlighted for clarity. The Bayesian posterior probability threshold is set at 0.90.

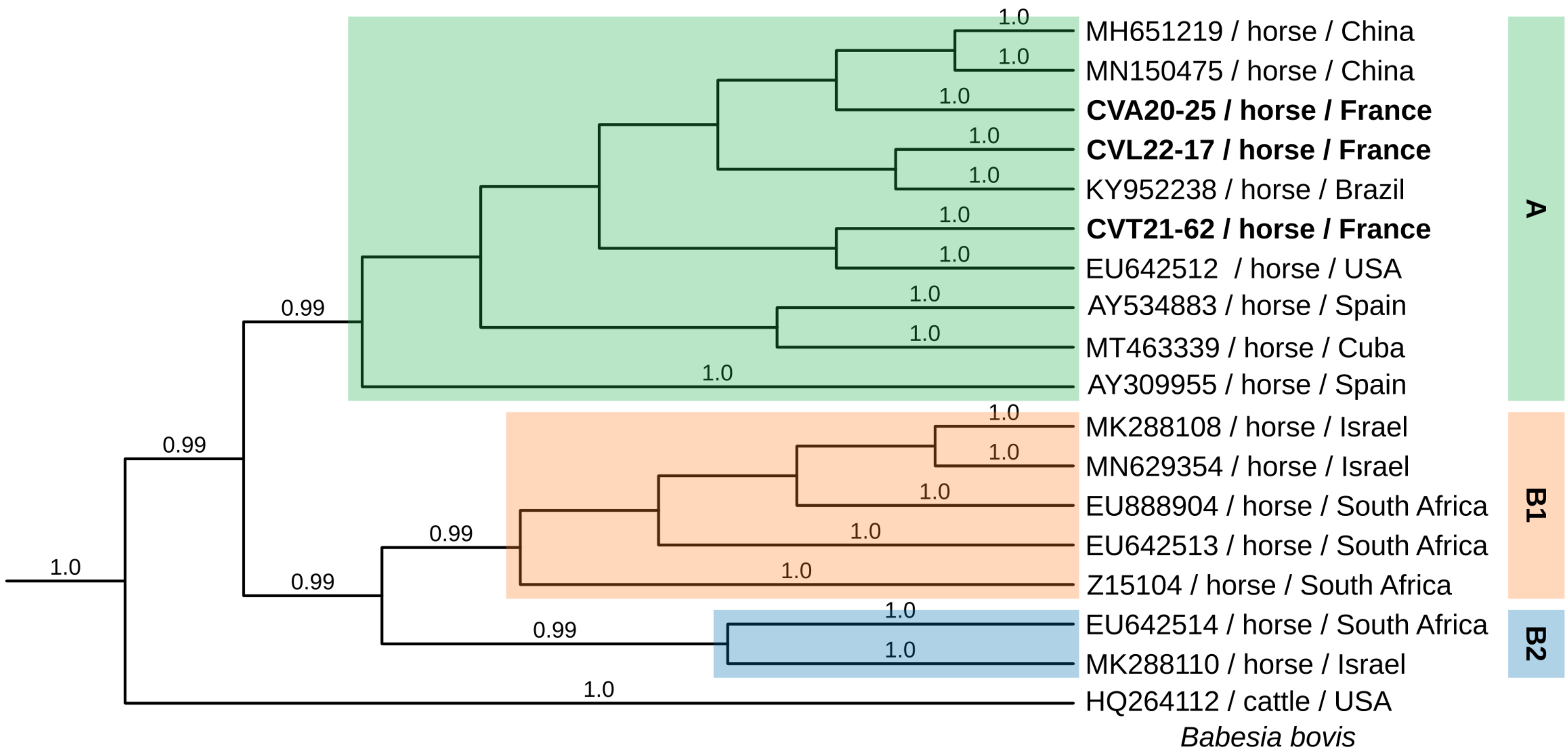


Supplementary Fig. 3. Phylogenetic tree of the 18S rRNA gene partial sequences of *Babesia caballi* constructed by Bayesian inference and represented as a cladogram. The tree comprises 38 sequences for which 1295 positions were analyzed. Sequences from this study are in bold and the tree genotypes (A, B1 and B2) have been highlighted for clarity. The Bayesian posterior probability threshold is set at 0.90.
